# “A distinct neutrophil population invades the central nervous system during pancreatic cancer”

**DOI:** 10.1101/659060

**Authors:** Kevin G. Burfeind, Xinxia Zhu, Mason A. Norgard, Peter R. Levasseur, Brennan Olson, Katherine A. Michaelis, Eileen Ruth S. Torres, Esha M. Patel, Sophia Jeng, Shannon McWeeney, Jacob Raber, Daniel L. Marks

**Affiliations:** Papé Family Pediatric Research Institute, Oregon Health & Science University, Portland, OR USA; Medical Scientist Training Program, Oregon Health & Science University, Portland, OR USA; Department of Behavioral Neuroscience, Oregon Health & Science University, Portland, OR USA; Division of Bioinformatics and Computational Biology, Department of Medical Informatics and Clinical Epidemiology, Oregon Health & Science University, Portland, OR USA; Oregon Clinical and Translational Research Institute, Oregon Health & Science University, Portland, OR USA; Knight Cancer Institute, Oregon Health & Science University, Portland, OR USA; Departments of Neurology and Radiation Medicine, Division of Neuroscience ONPRC, Oregon Health and & Science University, Portland, OR USA; Brenden-Colson Center for Pancreatic Care, Oregon Health and & Science University Portland, OR USA

**Keywords:** Neuroinflammation, Neuroimmunology, Neutrophils, Cancer Cachexia, Brain, Myeloid Cells

## Abstract

Weight loss, fatigue, and cognitive dysfunction are common symptoms in cancer patients that occur prior to initiation of cancer therapy. Inflammation in the brain is a driver of these symptoms, yet cellular sources of neuroinflammation during malignancy are unknown. In a mouse model of pancreatic ductal adenocarcinoma (PDAC), we observed early and robust myeloid cell infiltration into the brain. Infiltrating immune cells were predominately neutrophils, which accumulated at a unique central nervous system entry portal called the velum interpositum, where they expressed CCR2. CCR2 knockout mice had significantly decreased brain-infiltrating neutrophils as well as attenuated anorexia and muscle catabolism during PDAC, without any changes in neutrophils in other organs. Lastly, intracerebroventricular blockade of the purinergic receptor P2RX7 during PDAC abolished neutrophil recruitment to the brain and attenuated anorexia. Our data demonstrate a novel function for the CCR2/CCL2 axis in recruiting neutrophils to the brain, which drives anorexia and muscle catabolism.

## Introduction

Cancer patients commonly present with symptoms driven by disruption of normal CNS function. Weight loss, weakness, fatigue, and cognitive decline often occur in malignancies outside the CNS, and develop prior to initiation of cancer therapy (Meyers, 2000; Miller, Ancoli-Israel, Bower, Capuron, & Irwin, 2008). Many of these symptoms are part of a syndrome called cachexia, a devastating state of malnutrition characterized by decreased appetite, fatigue, adipose tissue loss, and muscle catabolism (Fearon et al., 2011). There are currently no effective treatments for cachexia or other CNS-mediated cancer symptoms. While mechanisms of CNS dysfunction during malignancy are still not well understood, inflammation in the brain is proposed as a key driver (Burfeind, Michaelis, & Marks, 2016). Inflammatory molecules (*e.g.*, lipopolysaccharide, cytokines) can cause dysfunction of the appetite-, cognition- weight-, and activity-regulating regions in the CNS, resulting in signs and symptoms nearly identical to those observed during cancer (T. P. Braun et al., 2011; Burfeind et al., 2016; Grossberg et al., 2011). Moreover, cytokines and chemokines are produced in these same regions during multiple types of cancer (Michaelis et al., 2017). Our lab and others previously showed that disrupting inflammatory signaling by deleting either MyD88 or TRIF attenuates anorexia, muscle catabolism, fatigue, and neuroinflammation during malignancy (Burfeind et al., 2018; Ruud et al., 2013).

The mechanisms by which inflammation generated in the periphery (*e.g.*, at the site of a malignancy) is translated into inflammation in the brain, and how this is subsequently translated CNS dysfunction, are still not known. Circulating immune cells present an intriguing cellular candidate, as they are thought to infiltrate and interact with the brain during various states of inflammation (Prinz & Priller, 2017), yet have not been investigated as potential mediators of brain dysfunction during cancer. Therefore, we characterized the identity, properties, and function of infiltrating immune cells in the brain using a syngeneic, immunocompetent, mouse model of pancreatic ductal adenocarcinoma (PDAC), a deadly malignancy associated with profound anorexia, fatigue, weakness, and cognitive dysfunction (Baekelandt et al., 2016; Michaelis et al., 2017). We observed that circulating myeloid cells, primarily neutrophils, were recruited to the CNS early in PDAC, infiltrating throughout the brain parenchyma and accumulating in the meninges near regions important for appetite, behavior, and body composition regulation. We then demonstrated a novel role for the CCR2-CCL2 axis (typically considered a monocyte chemotaxis pathway) in recruiting neutrophils specifically to the brain, rather than the liver or tumor. We also demonstrated that this axis is important for the development of anorexia and muscle catabolism during PDAC. Next, we blocked purinergic receptor P2RX7 signaling specifically on brain macrophages during PDAC via intracerebroventricular (ICV) injection of oxidized ATP (oATP), which prevented circulating myeloid cell recruitment to the brain and attenuated anorexia. Lastly, we showed that during PDAC, brain-infiltrating neutrophils have a transcriptional profile that is distinct from that of circulating, liver-infiltrating, and tumor-infiltrating neutrophils. Taken together, these results reveal a novel mechanism for neutrophil recruitment to the brain, in which a transcriptionally distinct population is recruited via an atypical neutrophil chemotactic factor, in a manner distinct from that in the periphery.

## Results

### Circulating myeloid cells infiltrate the brain early in PDAC

We first investigated whether circulating immune cells infiltrate the brain throughout the course of PDAC. We utilized a mouse model of PDAC, generated through a single intraperitoneal (IP) injection of C57BL/6 KRAS^G12D^ P53^R172H^ Pdx-Cre^+/+^ (KPC) cells. This well-characterized model recapitulates several key signs and symptoms of CNS dysfunction observed in humans, including anorexia, muscle catabolism, and fatigue (Michaelis et al., 2017). Using 10-color flow cytometry of whole brain homogenate (Fig. 1A), we characterized brain immune cells at three time points: 5 days post-inoculation (d.p.i) (before anorexia, fatigue, and muscle mass loss onset), 7 d.p.i. (initiation of wasting and anorexia), and 10 d.p.i. (robust wasting and anorexia, but 2-3 days before death) (see either ref 6 or Fig. 5F for typical disease progression in our KPC model). Compared to sham-injected animals, we observed a significant increase in CD45^high^CD11b+ myeloid cells in the brains of animals with PDAC (Fig. 1B), with an increase as a percentage of total CD45+ (all immune cells) and CD45^high^ (non-microglia leukocytes) cells occurring at 5 d.p.i. (Fig. 1D and Figure 1 – figure supplement 1D).

**Figure 1.**
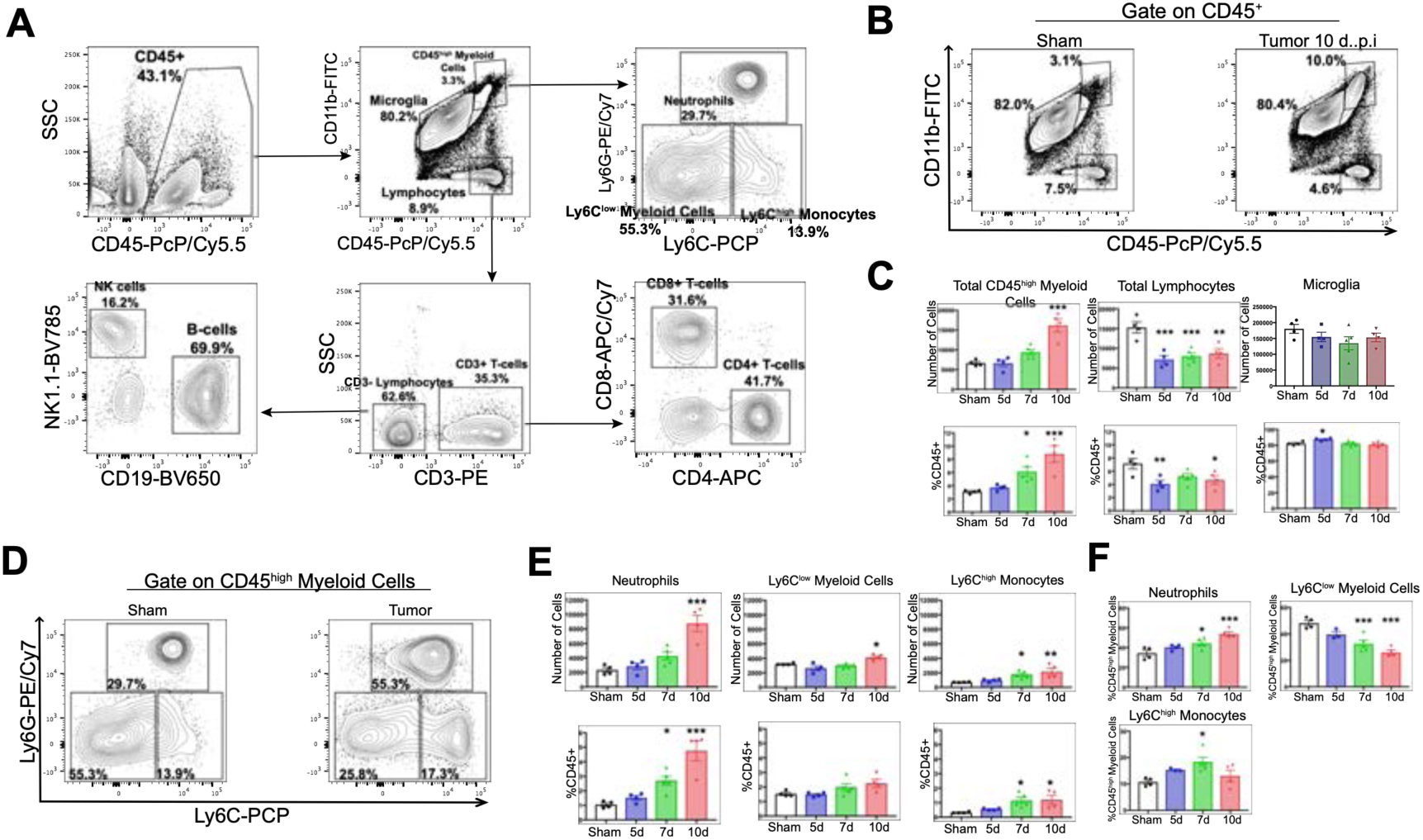
Myeloid cells infiltrate the brain during PDAC. A) Flow cytometry plots of immune cells isolated from whole brain homogenate, showing gating strategy to identify different immune cell populations. B) Representative flow cytometry plots displaying CD45 and CD11b fluorescent intensities of immune cells isolated from brains of tumor and sham animals, gated on live, singlet, CD45+ cells. C) Quantification of different immune cell populations in the brain at different time points throughout PDAC course. d = days post inoculation. Populations were identified as shown in Fig. 1a. D) Representative flow cytometry plots displaying Ly6C and Ly6G fluorescent intensities of immune cells isolated from brains of tumor and sham animals, gated on CD45^high^CD11b^high^ cells. E) Quantification of different CD45^high^ myeloid cell populations in the brain at different time points during PDAC progression. F) Relative amounts of different CD45^high^ myeloid cell populations as a percentage of total CD45^high^ myeloid cells, throughout the course of PDAC. Populations identified as described for Fig. 1e. *n* = 4-5/group, \**P* < 0.05, \*\**P* < 0.01, \*\*\**P* < 0.001 compared to sham group in one-way ANOVA Bonferroni *post hoc* analysis, and results are representative of three independent experiments. For all figures, data are presented as mean ± s.e.m.

Both absolute and relative number of lymphocytes (CD45^high^CD11b-) were decreased in the brains of tumor animals compared to sham animals starting at 5 d.p.i., which was driven by a decrease in B-cells and CD4+ T-cells (Fig. 1C and Figure 1 – figure supplement 1B-D). There was no change in number of microglia (defined as CD45^mid^CD11b+) throughout the disease course (Fig. 1C). Further phenotypic analysis of infiltrating myeloid cells revealed that by 7 d.p.i., there was an increase in relative number (as a percentage of total CD45+ and CD45^high^) of Ly6C^mid^Ly6G^high^ neutrophils, Ly6C^low^ myeloid cells, and Ly6C^high^ monocytes (Fig. 1D, E and Figure 1 – figure supplement 1D). We observed an increase in absolute number of neutrophils, Ly6C^high^ monocytes, and Ly6C^low^ myeloid cells starting at 7 d.p.i., which became significant at either 7 (Ly6C^high^ monocytes) or 10 d.p.i. (neutrophils and Ly6C^low^ myeloid cells) (Fig. 1D and E). Neutrophils were by far the most numerous invading myeloid cell type, constituting 34% percent of CD45^high^CD11b+ cells in sham animals, and increasing to nearly 54% by 10.d.p.i. in tumor animals (Fig. 1F).

Since the CD45^high^CD11b+ population we defined as invading circulating myeloid cells could also contain activated microglia, we generated GFP+ bone marrow chimera to determine if the majority of these cells were of peripheral origin. Furthermore, the population of CD45^high^CD11b+Ly6C^low^ myeloid cells could also be activated microglia. We generated GFP+ bone marrow chimera mice through conditioning WT mice with treosulfan to ablate marrow, then transplanting marrow from pan-GFP mice (Ly5.1^GFP^) (Figure 1 – figure supplement 2A). This system is advantageous because, unlike other alkylating agents, treosulfan does not cross or disrupt the blood brain barrier (Capotondo et al., 2012). On average, mice that underwent bone marrow transplant (GFP BMT mice) exhibited 75% chimerism (Figure 1 – figure supplement 2C). In agreement with results from WT marrow animals, we observed that at 10 d.p.i., thousands of GFP+ myeloid cells infiltrated the brain in tumor animals (Figure 1 – figure supplement 2B). The majority of these cells were neutrophils, with a concurrent increase in Ly6C^high^ monocytes (Figure 1 – figure supplement 2C-F). As we observed previously, this coincided with a decrease in brain lymphocytes (CD45+GFP+CD11b-) in tumor animals (Figure 1 – figure supplement 2D). We did not observe an increase in GFP+ Ly6C^low^ myeloid cells (Figure 1 – figure supplement 2F), suggesting that the increase in CD45^high^CD11b+Ly6C^low^ cells in our WT marrow PDAC mice was a result of microglia activation, rather than infiltrating monocytes.

Taken together, these data show that myeloid cells infiltrate the brain during PDAC, which correlates with symptom onset, and that there is a selective neutrophil recruitment. Since the purpose of this study was to investigate infiltrating cells, we chose to focus our subsequent analysis on myeloid cells, with an emphasis on neutrophils.

### Invading myeloid cells accumulate at CNS interfaces during PDAC

Prior studies demonstrated regional vulnerability in the CNS to immune cell invasion during systemic inflammation (D’Mello, Le, & Swain, 2009). Therefore, we investigated the anatomic distribution of infiltrating myeloid cells in the CNS. We performed immunofluorescence immunohistochemistry analysis at 10 d.p.i., since all tumor-inoculated animals reliably developed anorexia, fatigue, and muscle catabolism at this time point, yet were not at terminal stage (Michaelis et al., 2017). In addition, our flow cytometry analysis demonstrated a robust immune cell infiltrate in the brain at 10 d.p.i. For initial analysis, we defined leukocytes as CD45+ globoid cells. Although we observed scattered CD45+ globoid cells within the parenchyma in the cortex and thalamus in tumor mice (Figure 2 – figure supplement 1), we observed a robust increase in leukocytes in the meninges adjacent to the hippocampus and median eminence (ME) (Fig. 2B and C). We also quantified leukocytes in the area postrema, due to its importance for appetite and weight regulation (Fig. 2D). While there was an increase in overall CD45 immunoreactivity, these cells appeared ramified rather than globoid (60X inset in Fig. 2D), suggesting microglia activation rather than immune cell infiltration. We did not observe any CD45+ cells in the lateral parabrachial nucleus (data not shown), which was implicated in cancer-associated anorexia (Campos et al., 2017). This was perhaps due to its lack of proximity to a circumventricular organ or meninges. Interestingly, we observed an increase in neutrophils (defined as myeloperoxidase [MPO] positive, CD45+ globoid cells) only in the meninges surrounding the hippocampus (Fig. 2B). This layer of meninges, known as the velum interpositum (VI), is a double-layered invagination of the pia matter. This potential space is closed rostrally, communicates caudally with the quadrigeminal cistern, and is highly vascularized via a number of internal cerebral arterioles and veins. Recent studies demonstrate robust immune cell recruitment into the brain via this anatomical route after mild trauma, during CNS infection, and during CNS autoimmune disease (Alvarez & Teale, 2006; Schmitt, Strazielle, & Ghersi-Egea, 2012; Szmydynger-Chodobska, Shan, Thomasian, & Chodobski, 2016).

**Figure 2.**
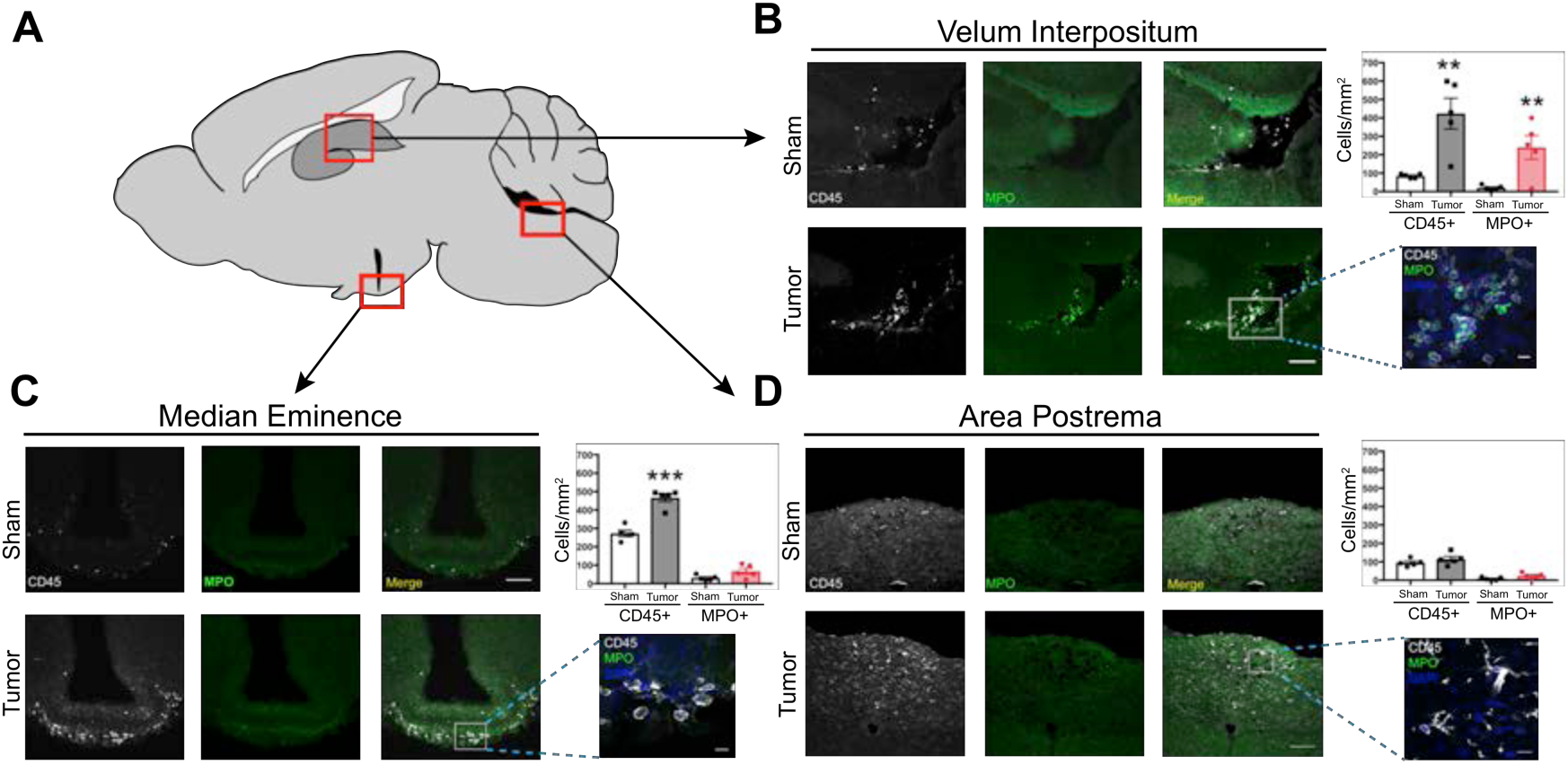
Infiltrating immune cells accumulate at CNS interfaces during PDAC cachexia. A) Picture of sagittal mouse brain section to illustrate different regions analyzed. B-) 20X images of velum interpositum (B), mediobasal hypothalamus (C), and area postrema (D) of brain from sham animal and tumor animal at 10 d.p.i., with 60X inset shown on the right, along with quantification of MPO+ and total CD45+ cells. Scale bar for 20X images = 100μm. Scale bar for 60X insets = 10μm. Data are presented as mean ± s.e.m., *n* = 5/group, \**P* < 0.05, \*\**P* < 0.01 in student’s t-test.

We verified the presence of meninges in the VI with ER-TR7 labeling, which showed infiltrating neutrophils in the VI meninges in tumor mice (Figure 2 – figure supplement 2A). Neutrophils in the VI were degranulating, with MPO “blebs” present at the edge of many cells, along with extracellular MPO (Figure 2 – figure supplement 2C). This phenomenon was only present in brains of tumor animals and not in brains of sham animals. We were able to confirm neutrophil identity with the plasma membrane marker Ly6G and globoid morphology (Figure 2 – figure supplement 1C). Neutrophil extracellular traps (NETs) were also present in the VI, as identified by citrillunated histone H3 and MPO co-labeling (Fig. 2G). We were unable to perform quantification on the number of NETs present in tumor mouse brains, due to the transient nature of these events.

In the CNS parenchyma, especially in the thalamus and cortex, we frequently observed neutrophils undergoing phagocytosis by microglia, with Iba-1+ cells extending processes around MPO+ neutrophils (Fig. 2H). This supports previous studies showing that microglia protect the CNS parenchyma from neutrophil invasion during various states of inflammation (Jens Neumann et al., 2018; J. Neumann et al., 2008; Otxoa-de-Amezaga et al., 2018).

The peripheral origin of the CD45+ globoid cells in the brain was assessed using our GFP BMT mice. Sham BMT mice showed very few GFP+ cells in the brain, including the cortex and thalamus (Fig. S2A), as well as the meninges (data not shown). In contrast, there was a large increase in GFP+ cells in the brains of KPC mice at 10 d.p.i. We observed a pattern of infiltrating GFP+ cells that was identical to CD45+ globoid cells in our previous experiments, with scattered GFP+ cells in the cortex and thalamus (Figure 2 – figure supplement 1A and B), and accumulations of GFP+ cells in the VI and ME (Figure 2 – figure supplement 2D and F). In agreement with our previous data, GFP+ cells were MPO+ in the VI (Fig. 2I), but not in the meninges of the ME (Figure 2 – figure supplement 2D). Most of the GFP+ cells in the ME were Iba-1+ (Figure 2 – figure supplement 2F), suggesting these cells were infiltrating monocytes that differentiated into meningeal macrophages. However, we did not observe any GFP+Iba-1+ cells in the CNS parenchyma.

Since we observed myeloid cell infiltration near brain regions important for behavioral and cognitive performance, we performed behavioral and cognitive tests on PDAC tumor mice to determine if these mice experienced altered anxiety levels or cognitive dysfunction (Figure 2 – Figure supplement 3). We observed that while PDAC tumor did not display cognitive dysfunction, they did spend significantly less time in the more anxiety-provoking center of the arena compared to sham mice, indicative of anxiety-like behavior, which confirms previous studies demonstrating anxiety during cancer (Campos et al., 2017) (Figure 2 – Figure supplement 3B-E).

### The CCR2-CCL2 axis is activated in the CNS during PDAC

Since *Ccl2* transcript showed the largest difference in induction between hippocampus and liver (17.1-fold vs. 3.6-fold) of all the chemokine transcripts we measured (with similar baseline levels of *Ccl2* expression, data not shown), and previous studies demonstrated the importance of CCR2/CCL2 for myeloid cell chemotaxis to the brain (Cazareth, Guyon, Heurteaux, Chabry, & Petit-Paitel, 2014; D’Mello et al., 2009), and PDAC cachexia in humans (Talbert et al., 2018), we chose to focus on the CCR2-CCL2 axis.

Using *in situ* hybridization we localized robust CCL2 mRNA expression exclusively within the VI during PDAC. There was no observable *Ccl2* mRNA in the brains of sham animals (Fig. 3B). We verified these results at the protein level using CCL2^mCherry^ mice, which showed abundant CCL2 protein expression in the VI in tumor animals at 10 d.p.i., exclusively expressed in Iba1+CD206+ meningeal macrophages. CCL2 protein was not expressed in VI meningeal macrophages in sham mice (Fig. 3C). We did not observe robust CCL2 protein expression in any other locations in the brain.

**Figure 3:**
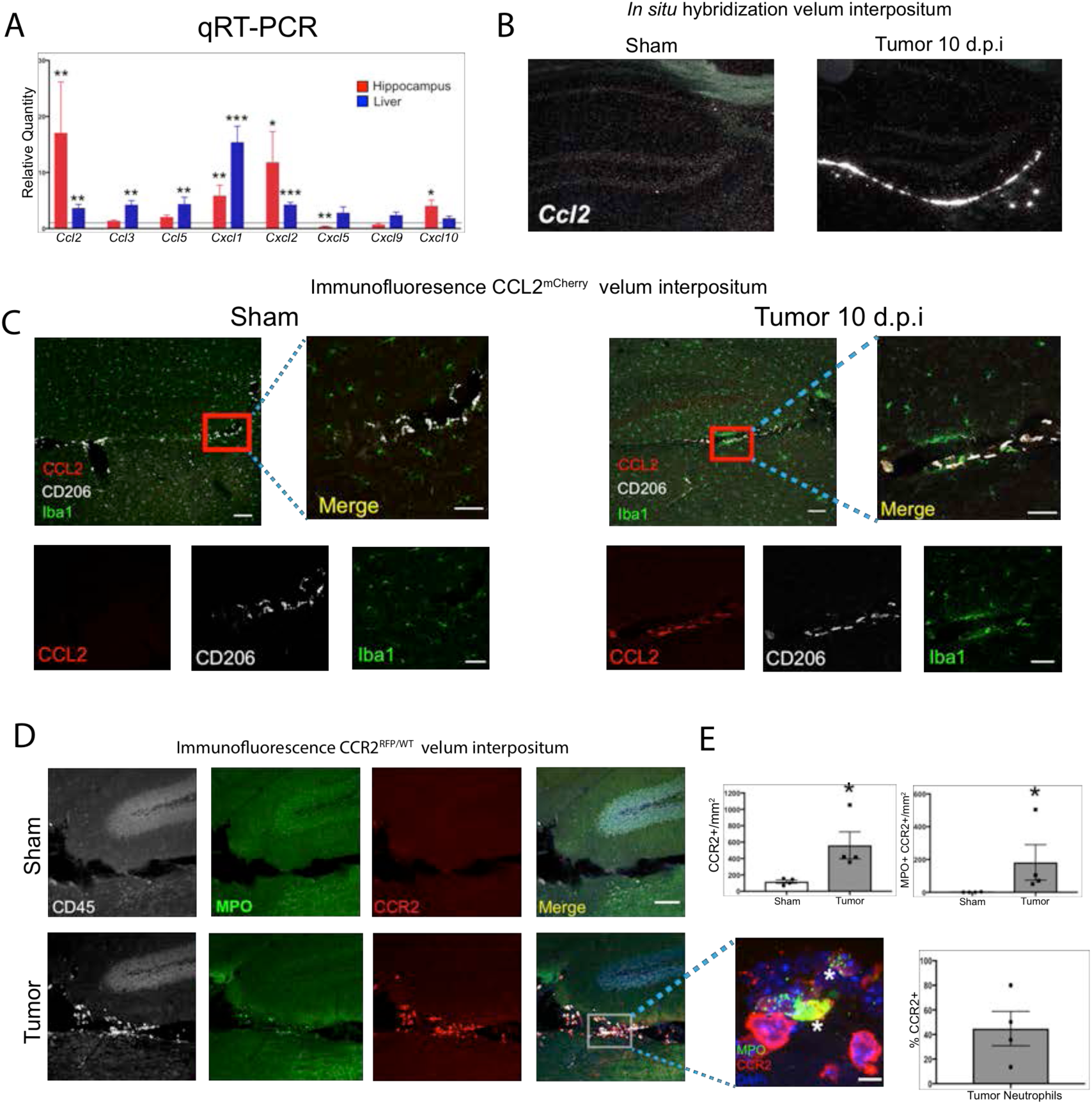
The CCR2-CCL2 axis is activated in the CNS during PDAC. A) qRT-PCR for select chemokine transcripts from dissected hippocampi and liver from tumor and sham animals 10 d.p.i. Values are tumor relative to sham. \**P* < 0.05, \*\**P* < 0.01, \*\*\**P* < 0.001 comparing tumor vs. sham within the same tissue type in student’s t-test. *n* = 5/group. Results are representative of two independent experiments. B) Representative darkfield microscopy image of *in situ* hybridization for CCL2 in sham and tumor (10 d.p.i) mouse brains. C) Representative 10X confocal microscopy image of brain from CCL2^mCherry^ tumor mouse brain, 10 d.p.i. Scale bar = 100 μm. Inset of VI shows CCL2 protein expression is confined to meningeal macrophages, identified by CD206 labeling. Scale bar = 20 μm. D) Representative 20X confocal microscopy image of brain from CCR2^RFP/WT^ tumor (10 d.p.i.) and sham mouse brain. Scale bar = 100 μm. Inset = 60X image identifying CCR2+ neutrophils in the VI of a tumor animal, indicated by asterisks. Scale bar = 5 μm. E) Quantification of different RFP+ cell populations in the VI of CCR2^RFP/WT^ tumor (10 d.p.i.) and sham animals. *n* = 4/group. *\*P* < 0.05, Mann-Whitney U-test comparing sham to tumor. Results are representative of two independent experiments.

CCR2^RFP/WT^ reporter mice were used to localize CCR2+ cells in the CNS. We observed that, at 10 d.p.i., CCR2+ immune cells infiltrated the brains of tumor mice and accumulated in the VI (Fig. 3D-F) Interestingly, a large percentage of neutrophils in the VI were CCR2+ (Fig. 3E), which infiltrated throughout the VI and often formed large aggregates consisting of 20 cells or more (Fig 3 – Figure Supplement 1A). CCR2+ cells were sparse or absent in other brain regions, particularly within the parenchyma, in tumor mice.

In order to verify CCR2 expression on neutrophils in the brains of tumor-bearing animals, we performed flow cytometry for CCR2 (using an anti-CCR2 antibody) on Ly6G+ circulating, liver-infiltrating, and brain-infiltrating neutrophils in both sham and PDAC-bearing animals at 10 d.p.i. As expected, we observed minimal CCR2 expression on circulating neutrophils in sham animals. While there was a slight increase in circulating CCR2+ neutrophils in tumor-bearing animals, there was no increase in CCR2+ neutrophils in the liver. Alternatively, there was a large increase in CCR2+ neutrophils in the brains of tumor-bearing animals (Fig 3 – Figure Supplement 1).

Based on our data showing that CCR2 is a brain-specific chemotactic receptor for neutrophils during PDAC, we hypothesized that brain-infiltrating neutrophils are unique compared to neutrophils that infiltrate other organs. In order to characterize the phenotype of brain-infiltrating neutrophils during PDAC, we performed RNA sequencing (RNAseq) on FACS-sorted neutrophils from blood, liver, tumor, and brain during PDAC at 10 d.p.i, as well as circulating neutrophils from sham animals (Figure 3 – Figure Supplement 2A and F). Principal component analysis of individual samples based on the top 500 most varying transcripts revealed that brain-infiltrating neutrophils clustered tightly together, but were distinct from those in liver, tumor, and blood (Figure 3 – Figure Supplement 2B). Furthermore, we were able to identify over 100 transcripts that were differentially expressed in the brain-infiltrating neutrophils compared to those in the liver, tumor, and circulation (Figure 3 – Figure Supplement 2C-E and Figure Supplement 3).

Taken together, these data indicate that the CCR2-CCL2 axis is activated in the CNS during PDAC, and that the CNS microenvironment uniquely influences the neutrophil transcriptome during PDAC.

### CCR2 is critical for neutrophil accumulation at CNS interfaces, anorexia, and muscle catabolism during PDAC

Based on our findings that CCR2+ cells infiltrate the brain during PDAC, we hypothesized that CCR2 is required for immune cell recruitment to the brain. We observed that at 11 d.p.i., there was a 37% decrease in total CD45^high^ myeloid cells in the brains of CCR2KO tumor mice compared to WT tumor mice (Fig. 4A and B). This difference was primarily driven by a large decrease in brain-infiltrating neutrophils in CCR2KO tumor mice, and to a much lesser extent a decrease brain-infiltrating Ly6C^high^ monocytes in CCR2KO tumor mice compared to WT tumor mice. There was a decrease in neutrophils (and again Ly6C^high^ monocytes, to a lesser extent) as a percentage of the CD45^high^ cells in the brain in CCR2KO tumor mice, indicating that the differences were not due to a global decrease in infiltrating immune cells (Fig. 4B). This was also supported by the fact that there were no differences in microglia (data not shown), Ly6C^low^ monocytes, or T-cells in the brains of CCR2KO tumor mice compared to WT tumor mice (Fig. 4C).

**Figure 4.**
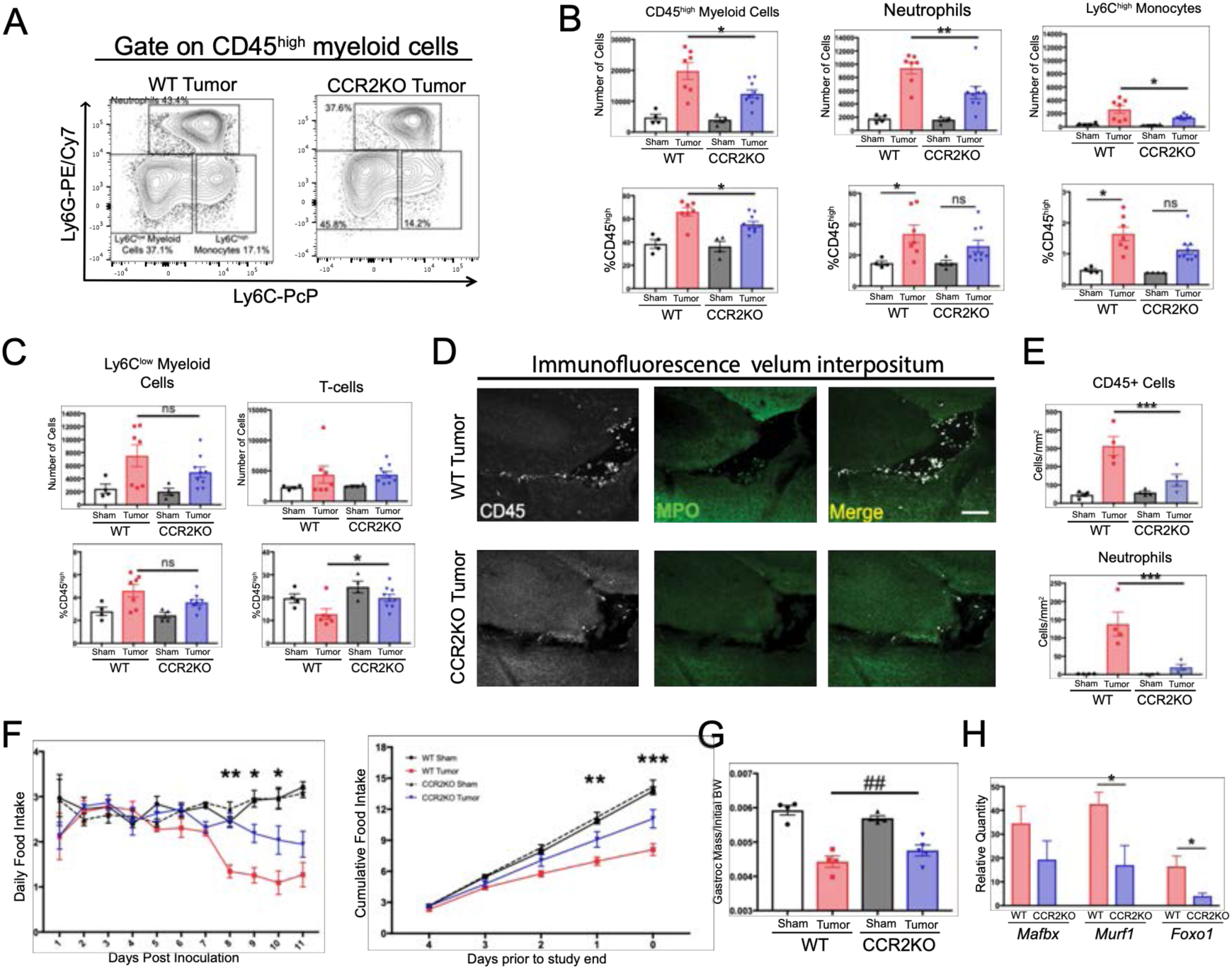
The CCR2-CCL2 axis in the CNS is critical for brain inflammation, anorexia, and muscle catabolism during PDAC. A) Representative plot of different CD45^high^ myeloid cell populations from WT and CCR2KO tumor animal brains, 11 d.p.i. Cells are gated on live, singlet, CD45+, CD45^high^CD11b+ cells. B and C) Flow cytometry analysis of immune cells isolated from whole brain homogenate. \**P* < 0.05, \*\**P* < 0.01, WT tumor vs. CCR2KO tumor, or tumor vs. sham in the same genotype in Bonferroni *post hoc* analysis in two-way ANOVA. ns = not significant. *n* = 4-9/group. Data consist of two independent experiments pooled (*n* = at least 2/group in each experiment). D) Representative 20X confocal microscopy images of the VI from WT tumor and CCR2KO tumor brain, 10 d.p.i. Scale bar = 100 μm. E) Quantification of total CD45+ globoid cells and MPO+ neutrophils in the VI of WT and CCR2KO tumor and sham animals, 10 d.p.i. \*\*\**P* < 0.001, WT tumor vs. CCR2KO tumor in Bonferroni *post hoc* analysis in two-way ANOVA. *n* =4/group. F) Daily food intake (left) and final 5 days of the study (right, starting when animals develop symptoms) in WT and CCR2KO tumor and sham mice. \**P* < 0.05, \*\**P* < 0.01, \*\*\**P* < 0.001 comparing WT tumor vs. CCR2KO tumor in Bonferroni *post hoc* analysis in two-way ANOVA. *n* = 4/5 per group. Results are representative of three independent experiments. G) Left = mass of dissected gastrocnemius, normalized to initial body weight, at 11 d.p.i. ##*P* < 0.01 for interaction effect between genotype and tumor status in two-Way ANOVA analysis. H) qRT-PCR analysis of *Mafbx*, *Murf1*, and *Foxo1* from RNA extracted from gastrocnemii dissected at 11 d.p.i. Values normalized to those from WT sham. \**P* < 0.05, WT tumor vs. CCR2KO tumor dCt values. *n* = 3-5/group.

**Figure 5.**
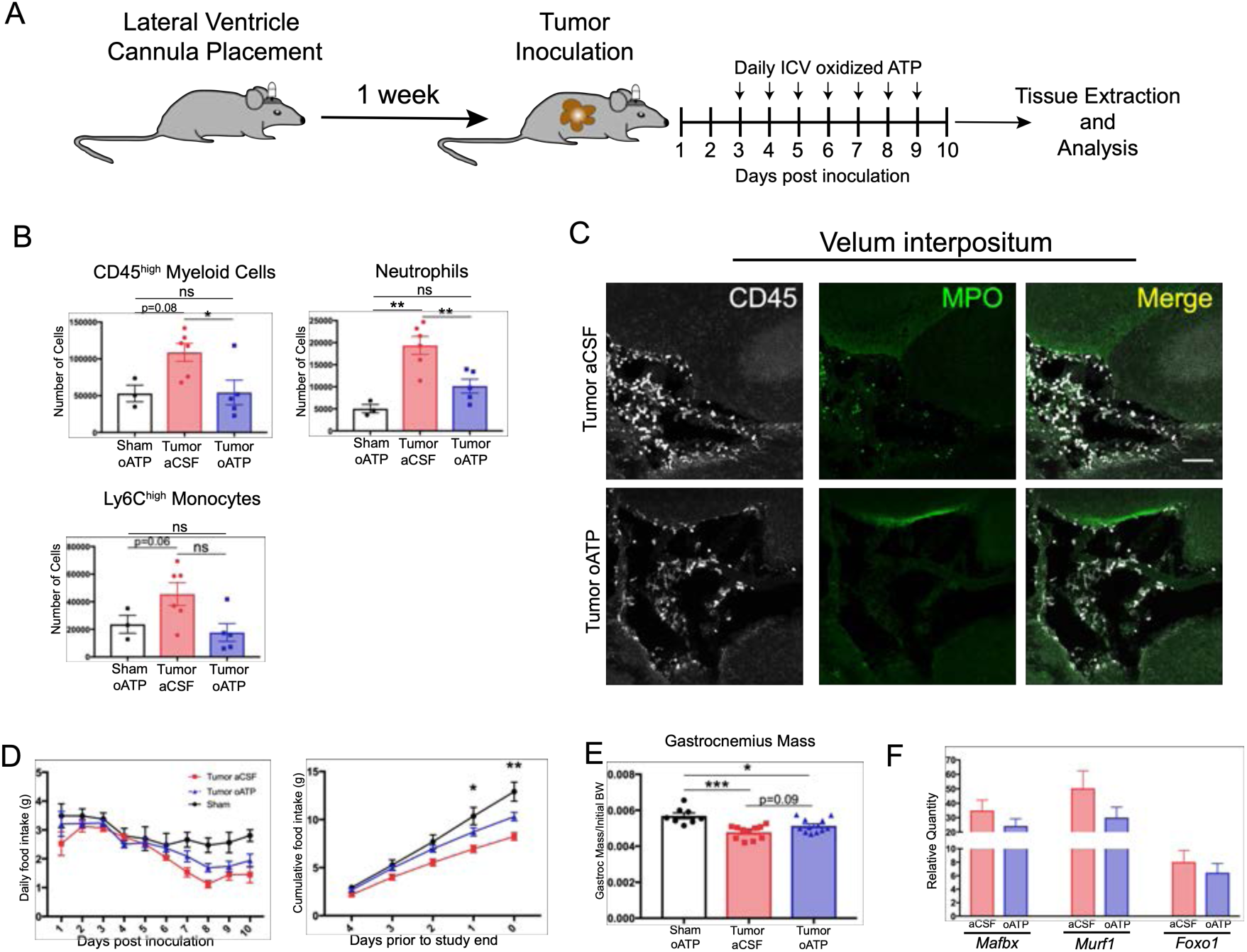
Intracerebroventricular administration of oxidized ATP prevents immune cell recruitment to the brain and attenuates anorexia during PDAC. A) Diagram depicting workflow for lateral ventricle cannulation and ICV oATP treatment during PDAC. ICV = intracerebroventricular. B) Quantification of immune cells isolated from whole brain homogenate. \**P* < 0.05, \*\**P* < 0.01, in Bonferroni *post hoc* analysis in two-way ANOVA. ns = not significant. *n* = 4-7/group. C) Representative 20X confocal microscopy images of the VI from aCSF-treated and oATP-treated tumor animals. Scale bar = 100 µm. D) Daily food intake (left) and cumulative food intake for the final 5 days of the study (right, starting when animals develop symptoms) \**P* < 0.05, \*\**P* < 0.01, comparing aCSF tumor vs. oATP tumor in Bonferroni *post hoc* analysis in two-way ANOVA. *n* = 8-12/group. Results consist of two independent experiments pooled (*n* = 4-7/group in each experiment). E) Left = mass of dissected gastrocnemius, normalized to initial body weight, at 10 d.p.i. F) qRT-PCR analysis of *Mafbx*, *Murf1*, and *Foxo1* from RNA extracted from gastrocnemii dissected at 8-10 d.p.i. Values normalized to those from sham oATP. *n* = 4-7/group.

Since CCR2+ immune cells, particularly neutrophils, in the brains of tumor mice localized primarily to the VI, we hypothesized that there would be a decrease in immune cells in the VI in CCR2KO tumor animals. Indeed, we observed a dramatic decrease in both total CD45+ globoid and MPO+ immune cells in the VI in CCR2KO tumor mice compared to WT tumor mice (Fig. 4D and E).

We previously demonstrated that KPC-bearing animals experienced robust anorexia and muscle catabolism (Michaelis et al., 2017), which our lab and others showed are driven by CNS inflammation (T. P. Braun et al., 2011; Laye et al., 2000). We observed that CCR2 knockout (CCR2KO) mice had decreased anorexia during PDAC compared to WT tumor mice (Fig. 4A). CCR2KO tumor mice also had attenuated muscle loss compared to WT tumor mice (Fig. 6B). To determine whether the decreased muscle mass loss in CCR2KO mice was due to decreased muscle proteolysis, we assessed levels of transcripts key for muscle proteolysis in the gastrocnemius, including *Mafbx*, *Murf1*, and *Foxo1*, which we previously showed are induced by CNS inflammation (T. P. Braun et al., 2011). We observed that, compared to WT tumor animals, CCR2KO tumor animals had decreased induction of *Murf1* and *Foxo1* (Fig. 4C), confirming that there was decreased muscle catabolism in CCR2KO tumor mice.

Since CCR2 deletion was not brain specific in the CCR2KO mice, we performed an extensive analysis of infiltrating immune cells in other organs (Fig. 4 – Figure Supplement 1). We observed minimal changes in immune cell composition in the blood, liver, and tumor in CCR2KO tumor mice compared to WT tumor mice. We only observed a decrease in circulating Ly6C^high^ monocytes in CCR2KO tumor mice (Fig. 4 – Figure Supplement 1G). When we assessed neutrophils in CCR2KO tumor mice in different organs as a percentage of those in WT tumor mice, we found the largest decrease to be in the brain and observed a slight increase in circulating neutrophils in CCR2KO tumor mice compared to WT tumor mice (Fig. 4 – Figure Supplement 1F and H), suggesting that the decrease in brain-infiltrating neutrophils was due to a homing defect rather than inability to mobilize from the bone marrow. Therefore, our data show that CCR2 is important for neutrophil recruitment specifically to the brain, and that the decrease in brain-infiltrating neutrophils was due to a homing defect rather than inability to mobilize from the bone marrow.

Lastly, although there was a 47% decrease in brain-infiltrating Ly6C^high^ monocytes in CCR2KO tumor mice compared to WT tumor mice, we also observed a 59% decrease in circulating Ly6C^high^ monocytes, suggesting that, unlike neutrophils, the decrease in brain-infiltrating Ly6C^high^ monocytes was in fact due to a defect in marrow extravasation.

Taken together, these data show that the CCR2-CCL2 axis required for myeloid cell recruitment specifically to the CNS during PDAC, and that this axis is important for development of anorexia and muscle catabolism.

### Blockade of P2RX7 in the CNS prevents immune cell infiltration into the brain and attenuates cachexia during PDAC

To evaluate the effects of CNS inflammatory responses during PDAC independent of potential systemic effects, we treated mice with intracerebroventricular (ICV) oxidized ATP (oATP). This potently blocks purinergic receptor P2RX7 signaling on brain resident macrophages. Signaling through this receptor is key for neutrophil recruitment to the brain during neuroinflammation (Roth et al., 2014). Animals were surgically implanted with indwelling lateral ventricle cannulas, then inoculated with KPC cells one week later. Mice received daily ICV injections of either 500 ng oATP or vehicle (aCSF), starting 3 d.p.i. (Fig. 5A). oATP treatment completely prevented both neutrophils and total CD45^high^ myeloid cells from infiltrating the brain (Fig. 5B). Ly6C^low^ myeloid cells and T-cells were not affected (Figure 5 – Figure supplement 1B). Furthermore, ICV oATP treatment did not affect any circulating immune cell population (Figure 5 – Figure supplement 1C). When we investigated infiltrating immune cells in the VI, both CD45+ globoid cells and CD45+MPO+ neutrophils were completely absent in oATP-treated tumor animals, compared to large infiltrates in aCSF-treated tumor animals (Fig. 5C). While we did observe sparse CD45+ cells in the VI in oATP-treated tumor animals, they were not globoid and resembled meningeal macrophages. We also observed that oATP treatment attenuated anorexia in tumor mice (Fig. 5D). There was trend toward increased gastrocnemius mass (*P* = 0.09) in oATP-treated tumor mice compared to aCSF-treated tumor bearing mice (Fig. 5E), which corresponded to a trend toward decreased induction of genes associated with proteolysis in gastrocnemius muscle (Fig. 5F), demonstrating that muscle catabolism was moderately attenuated by oATP administration directly into the brain. Tumor size in oATP-treated tumor mice was identical to that of aCSF-treated tumor mice (Figure 5 – Figure supplement 1A).

Since ICV oATP antagonizes P2RX7 on brain macrophages, we investigated its effect on microglia. To quantify activation state we assessed microglia morphology in the hippocampus. We did not observe any differences in microglia size, Iba-1 staining area, and Iba-1 intensity per cell when comparing aCSF- or oATP-treated tumor animals to oATP-treated sham animals or each other (Figure 5 – Figure supplement 2). These results show that microglia activation state in the hippocampus is not affected by the presence of a pancreatic tumor or oATP administration to the brain.

## Discussion

Several lines of investigation show that production of inflammatory mediators in the brain correlates strongly with CNS-mediated symptoms during cancer (Burfeind et al., 2018; Michaelis et al., 2017), yet the role of neuroinflammation during malignancies outside the CNS is still not well understood. Our data show that in a mouse model of PDAC, myeloid cells, consisting predominately of neutrophils, infiltrate the brain, in a CCR2-dependent manner, where they drive anorexia and muscle catabolism. We observed that infiltrating immune cells accumulated specifically in a unique layer of meninges called the velum interpositum (VI), which is adjacent to the hippocampus and the habenula, the latter of which is important for appetite regulation and is associating with cachexia in humans (Maldonado et al., 2018). We observed robust CCL2 mRNA and protein expression, along with CCR2+ neutrophils, exclusively in this region. The VI is implicated as a key structure for initial immune infiltration during states of neuroinflammation such as EAE (Schmitt et al., 2012) and traumatic brain injury (Szmydynger-Chodobska et al., 2016). Indeed, the VI contains the pial microvessels that are a key aspect of the “gateway reflex”, a neuro-immune pathway that involves interactions between leukocytes and neurons involved in stress response (Tanaka, Arima, Kamimura, & Murakami, 2017) and is implicated in gastrointestinal dysfunction during EAE (Arima et al., 2017). While we observed myeloid cell infiltration throughout the VI, we also observed accumulation of neutrophils and other leukocytes around the same pial vessels involved in the gateway reflex. The role of the gateway reflex in feeding behavior has not been investigated. It is possible that, in our model of PDAC, brain infiltrating neutrophils were involved in generating anorexia and muscle catabolism via a neuro-immune circuit similar to the gateway reflex, involving inflammation generated in the VI, and possibly transmitted to the habenula, or other regions involved in appetite regulation.

The role and presence of infiltrating leukocytes in the CNS during systemic inflammation remain poorly understood. While previous reports show that neutrophils infiltrate the brain after septic doses of LPS or sepsis induced by cecal ligation (He et al., 2016), it is still unknown if they contribute to neurologic sequelae (anorexia, fatigue, cognition and memory deficits, etc.) during and after sepsis. A series of studies utilizing a mouse model of inflammatory liver disease showed that “sickness behaviors” could be attenuated if myeloid cell recruitment to the brain was abrogated via any one of several different interventions, including: 1) administration of a P-selectin inhibitor (Kerfoot et al., 2006), 2) deleting CCR2 (D’Mello et al., 2009), and 3) inhibiting microglia activation with minocycline (D’Mello et al., 2013). However, unlike our study, these studies did not address many CNS-mediated signs and symptoms associated with chronic disease, including anorexia and muscle catabolism, instead using social interaction as their sole measure of sickness behaviors. They also did not address whether their interventions affected monocyte infiltration in other tissues. In addition to systemic inflammatory diseases, Zenaro et al. showed that transient neutrophil depletion led to substantially improved amyloid beta burden, decreased neuroinflammation, and lessened cognitive decline in a mouse model of Alzheimer’s disease (Zenaro et al., 2015). Therefore, our results, along with previous studies, implicate brain-infiltrating myeloid cells as key players in driving CNS-mediated signs and symptoms during inflammatory disease.

We observed a decrease in total number of lymphocytes in the brain starting at 5 d.p.i., which persisted throughout the course of PDAC. This was driven by a decrease in B-cells and CD4+ T-cells. Since the vast majority of lymphocytes in the non-inflamed murine brain are intravascular, even after thorough perfusion of the vasculature (Mrdjen et al., 2018), we chose not to pursue lymphocytes in our subsequent analysis. However, these interesting results warrant investigation of the role of intravascular lymphocytes in the brain during conditions of inflammation. While several studies showed that intravascular neutrophils can induce pathology in the brain (Atangana et al., 2017; Ruhnau, Schulze, Dressel, & Vogelgesang, 2017), the function of intravascular T-cells, B-cells, and NK cells, which are presumably adherent to the endothelium, has yet to be investigated.

We performed the open field test and object recognition test to assess behavioral alterations and cognitive injury in KPC-derived tumor mice. We observed that tumor-bearing mice spent significantly less time in the center of the arena during the open field test, indicative of anxiety-like behavior. Our results are consistent with previous studies on animals inoculated with Lewis lung carcinoma cells, showing that tumor animals display anxiety-like behaviors (Campos et al., 2017; McGinnis et al., 2017). However, it is important to note that tumor animals moved significantly less than sham animals. Therefore, the severe decrease in locomotor activity animals experienced may affect the ability to detect alterations in anxiety-like behavior and complicates extensive behavioral analysis in this model. The preferential exploring of the novel object in the object recognition test indicates that in contrast to activity, cognition was not impaired in this model.

We showed that CCR2KO mice exhibited significantly attenuated myeloid cell infiltration into the brain, as well as decreased anorexia and muscle catabolism, during PDAC. As discussed above, these results are in agreement with previous studies investigating sickness behaviors during inflammatory liver disease, which showed that CCR2KO mice exhibited attenuated monocyte infiltration into the brain, along with decreased sickness behaviors (D’Mello et al., 2009). Furthermore, it was recently reported that mice lacking CCR2 had decreased myeloid cell infiltration into the brain and attenuated cognitive impairment during a model of sepsis induced by *Streptococcus pneumoniae* injection into the lungs (Andonegui et al., 2018). In an attempt to identify inflammatory biomarkers for PDAC-associated cachexia, Talbert et al. identified CCL2 as the only cytokine or chemokine (out of a panel of 25) that was increased in the serum of cachectic PDAC patients but not increased in the serum of non-cachectic patients (Talbert et al., 2018). It is possible that the differences we observed in gastrocnemius catabolism between WT and CCR2KO tumor animals were due to differences in food intake, but the fact that we observed a significant decrease in induction of the catabolic genes *Mafbx*, *Murf1*, and *Foxo1* in CCR2KO tumor animals, which are not induced by decreased food intake/starvation (T. P. Braun et al., 2011), makes this unlikely.

While CCR2 is usually not considered a key receptor for neutrophil recruitment, previous studies show it is important for neutrophil chemotaxis during sepsis (Souto et al., 2009; Souto et al., 2011). Interestingly, while we observed a robust decrease in brain-infiltrating neutrophils, we did not observe a decrease in liver- or tumor-infiltrating neutrophils in CCR2KO tumor mice, indicating that CCR2 is important for neutrophil recruitment specifically to the brain. Circulating neutrophils in sham animals did not express CCR2, but a small percentage of circulating neutrophils expressed CCR2 in tumor animals. There was no increase in CCR2-expressing neutrophils in the liver during PDAC. Alternatively, a significant percentage of neutrophils in the brain expressed CCR2 during PDAC, meaning a distinct population of neutrophils is recruited to the brain from the circulation. In addition, there was actually a small increase in circulating neutrophils in CCR2KO tumor mice, ruling out the possibility that neutrophils were unable to extravasate out of the marrow. These results, along with our RNASeq data (discussed below), suggest that the population recruited to the brain has a distinct function from those recruited to other organs.

We administered oxidized ATP, a purinergic receptor antagonist, directly into the brain and observed complete abrogation of circulating myeloid cell recruitment to the brain in tumor animals, as well as anorexia attenuation. These results provide key mechanistic insights to show that brain inflammation is key for PDAC-associated anorexia. While there was no change in microglia morphology after oATP administration, consistent with previous studies (Martin et al., 2018; Roth et al., 2014), we cannot rule out the possibility that the difference in anorexia we observed were due to changes in microglia phenotype. The presence of an indwelling lateral ventricle cannula may have also induced microglia activation and influenced morphology quantification. However, we did take care to acquire images from the contralateral hemisphere. Furthermore, we observed an increased Ly6C^high^ monocyte infiltrate in our aCSF-treated tumor animals compared to non-cannulated tumor animals, suggesting the indwelling lateral ventricle cannula did affect the inflammatory response in the brain to at least a small degree. Nevertheless, oATP completely prevented myeloid cells from infiltrating the brain during PDAC, strongly implicating these cells as mediators of anorexia.

A few limitations should be considered when interpreting results of this study. First, our data were produced in a single model of pancreatic cancer. While our model is extensively characterized and reliably recapitulates many of the CNS-mediated symptoms observed in humans, other malignancies should also be considered. Second, it is possible, even likely, that circulating immune cells infiltrate and influence function in other organs dysfunctional during cancer (skeletal muscle, adipose tissue, etc.). However, the purpose of this study was to investigate and characterize interactions between circulating immune cells and the brain during PDAC. Therefore, we chose to focus specifically on the brain so as to not overcomplicate analysis. Third, we did observe a small increase in Ly6C^hi^ monocytes in the brains of animals during PDAC, which was attenuated by CCR2 deletion. We cannot rule out that these cells did not contribute to anorexia and muscle catabolism. However, there were far fewer Ly6C^hi^ monocytes (≈2,000) in the brain than neutrophils (≈9,000) during PDAC, and these cells only constituted about 15% of brain CD45 high myeloid cells (vs. about 50% for neutrophils). Lastly, despite our extensive analysis, we cannot rule out with absolute certainty that the differences we observed in CCR2KO mice were not due to differences in tumor response. However, both the CCR2/CCL2 axis and neutrophils are reported to be “pro-tumor” (Coffelt, Wellenstein, & de Visser, 2016; Qian et al., 2011) and therefore systemic treatment targeting neutrophils or the CCR2/CCL2 axis in humans may be particularly beneficial in that they decrease tumor size and abrogate CNS dysfunction. This would be advantageous to conventional anti-tumor therapies such as chemotherapy and checkpoint inhibitors, which are both known to cause cachexia-like symptoms (Theodore P. Braun et al., 2014; Michot et al., 2016) and not effective against PDAC (especially checkpoint inhibitors).

In summary, we demonstrated that myeloid cells infiltrate the CNS throughout the course of PDAC and that preventing myeloid cells from infiltrating the brain attenuates anorexia and muscle catabolism. We showed there are distinct mechanisms for immune cell recruitment to the brain during systemic inflammation, and demonstrate a novel role for CCR2 in neutrophil recruitment to the brain, providing key insights into mechanisms of neuroinflammation and associated symptoms.

## Materials and Methods

### Animals

Male and female 20–25g WT C57BL/6J (stock no. 000664), Ly5.1-EGFP (stock no. 00657), CCL2^mCherry^ (stock no. 016849), and CCR2KO (stock no. 004999) were purchased at Jackson Laboratories. Animals were aged between 7 and 12 weeks at the time of study and maintained at 27°C on a normal 12:12 hr light/dark cycle and provided *ad libitum* access to water and food. Experiments were conducted in accordance with the National Institutes of Health Guide for the Care and Use of Laboratory Animals, and approved by the Animal Care and Use Committee of Oregon Health & Science University.

### KPC Cancer Model

Our lab generated a mouse model of PDAC by a single IP injection of murine-derived KPC PDAC cells (originally provided by Dr. Elizabeth Jaffee from Johns Hopkins) (Michaelis et al., 2017). These cells are derived from tumors in C57BL/6 mice heterozygous for oncogenic <u>K</u>RAS^G12D^ and point mutant T<u>P</u>53^R172H^ with expression targeted to the pancreas via the PDX-1-<u>C</u>re driver (Foley et al., 2015). Cells were maintained in RPMI supplemented with 10% heat-inactivated FBS, and 50 U/mL penicillin/streptomycin (Gibco, Thermofisher), in incubators maintained at 37°C and 5% CO_2_. In the week prior to tumor implantation, animals were transitioned to individual housing to acclimate to experimental conditions. Animal food intake and body weight were measured once daily. Sham-operated animals received PBS in the same volume. Bedding was sifted daily to account for food spillage not captured by cagetop food intake measurement. Animals were euthanized between 8 and 11 days post inoculation, when food intake was consistently decreased and locomotor activity was visibly reduced, yet signs of end-stage disease (ascites, unkempt fur, hypotheremia, etc.) were not present (Michaelis et al., 2017).

### Generation of Ly5.1-EGFP Chimera Mice

WT C57BL/6J male mice aged 8-10 weeks were injected IP with the alkylating agent treosulfan (Ovastat^®^, a generous gift from Joachim Baumgart at Medac GmbH, Germany) at a dose of 1500 mg/kg/day for 3 consecutive days prior to the day of bone marrow transplant (BMT). 24 hrs after the third treosulfan injection, a Ly5.1-EGFP male or female donor mouse aged between 2-6 months was euthanized and femurs, tibias, humeri, and radii were dissected. After muscle and connective tissue were removed, marrow cells were harvested by flushing the marrow cavity of dissected bones using a 25-gauge needle with Iscove’s modified Dulbecco’s medium supplemented with 10% FBS. The harvested cells were treated with RBC lysis buffer, filtered with a 70 μm cell strainer, and counted. 3-4 × 10^6^ cells in 200 μL HBSS were transplanted immediately into each recipient mouse via tail vein injection. To prevent infection during an immunocompromised period, recipient mice received amoxicillin dissolved in their drinking water (150 mg/L) for 2 weeks starting on the first day of treosulfan injection. GFP BMT mice were given at least 5 weeks for marrow reconstitution and recovery. Percent chimerism in each GFP BMT mouse was determined by flow cytometry analysis of circulating leukocytes.

### Behavioral Analysis

Behavioral and cognitive tests were performed on days 7-9 d.p.i. The open field test was conducted on days 7 and 8 post-inoculation, and the object recognition test was performed on days 8 and 9 post-inoculation. For all behavioral analyses, observers were blinded to group (tumor vs. sham).

#### Open field Testing

Exploratory and anxiety-like behaviors were assessed using the open field test on two subsequent days. The open field consisted of a brightly lit square arena (L 40.6 × W 40.6 × H 40.6 cm). The light intensity in the center of the open field was 100 lux. Mice were allowed to explore for 10 min in each trial. Behavioral performance was tracked and analyzed using an automated video system (Ethovision 7.0 XT, Noldus). Exploratory behavior was analyzed and included total distance moved and time spent in the center (20 × 20 cm) of the open field.

#### Novel Object Recognition

Mice were habituated to the open field arena over two days as described above on two subsequent days. On the third day, mice were exposed to the arena containing two identical objects (small orange hexagonal prisms) placed 15 cm from the adjacent walls and 10 cm apart for 15 min. On day four, one of the identical objects (“familiar”) was replaced with a novel object (small green triangular prism) of similar dimensions and mice were again allowed to explore for 15 min. During both the open field and novel object recognition tests, mice were placed into the center of the arena. Clear visuospatial orientation to the object, within 2 cm proximity, as well as physical interaction with the object was coded as exploratory behavior, and the percent time spent exploring the novel versus the familiar object was calculated. Three tumor animals were excluded from analysis because of complete lack of exploratory behavior.

### Intracerebroventricular Cannulation and Injections

Mice were anesthetized under isoflurane and placed on a stereotactic alignment instrument (Kopf Instruments). 26-gauge lateral ventricle cannulas were placed at 1.0 mm X, −0.5 mm Y, and −2.25 mm Z relative to bregma. Mice were given one week for recovery after cannula placement. Injections were given in 2 µl total volume. Oxidized ATP was dissolved in aCSF and injected at a concentration of 250 ng/μL over 5 min while mice were anesthetized under isoflurane.

### Immunofluorescence Immunohistochemistry

Mice were anesthetized using a ketamine/xylazine/acetapromide cocktail and sacrificed by transcardial perfusion fixation with 15 mL ice cold 0.01 M PBS followed by 25 mL 4% paraformaldehyde (PFA) in 0.01 M PBS. Brains were post-fixed in 4% PFA overnight at 4°C and cryoprotected in 20% sucrose for 24 hrs at 4°C before being stored at −80°C until used for immunohistochemistry. Immunofluorescence immunohistochemistry was performed as described below. Free-floating sections were cut at 30 μm from perfused brains using a Leica sliding microtome. Sections were incubated for 30 min at room temperature in blocking reagent (5% normal donkey serum in 0.01 M PBS and 0.3% Triton X-100). After the initial blocking step, sections were incubated in primary antibody (listed below) in blocking reagent for 24 hrs at 4°C, followed by incubation in secondary antibody (also listed below) for 2 hrs at room temperature. Between each stage, sections were washed thoroughly with 0.01 M PBS. Sections were mounted onto gelatin-coated slides and coverslipped using Prolong Gold antifade media with DAPI (Thermofisher).

The following primary anti-mouse antibodies were used, with company, clone, host species, and concentration indicated in parentheses: CD11b (eBioscience, rat, M1/70, 1:1000), CD45 (BD, rat, 30-F11, 1:1000), myeloperoxidase (R&D, goat, polyclonal, 1:1000), Ly6G (Biolegend, 1A8, rat, 1:250), Iba-1 (Wako, Rabbit, NCNP24, 1:1000), CD206 (Bio-rad, rat, MR5D3, 1:1000), ER-TR7 (Abcam, rat, ER-TR7, 1:1000), and citrillunated histone H3 (Abcam, rat, polyclonal, 1:1000). We also used a chicken anti-mCherry antibody (Novus Biologicals, polyclonal, 1:20,000), to amplify mCherry signal in sections from CCL2^fl/fl^ mice and a rabbit anti-RFP antibody (Abcam, polyclonal, 1:1000) to amplify RFP signal in sections from CCR2^RFP/WT^ mice.

The following secondary antibodies were used, all derived from donkey and purchased from Invitrogen, with dilution in parentheses: anti-goat AF488 (1:500), anti-rabbit AF555 (1:1000), anti-rat AF555 (1:1000), anti-rat AF633 (1:500), and anti-chicken AF555 (1:1000).

### Image acquisition and analysis

All images were acquired using a Nikon confocal microscope. Cell quantification was performed on 20X images using the Fiji Cell Counter plugin by a blinded researcher. CD45+ cells were defined as CD45 bright globoid cells, and neutrophils were defined as CD45+ MPO+ cells. The velum interpositum (VI) was defined as the layer of meninges (identified by appearance of staining background) between the hippocampus and thalamus, from bregma −1.7 to −2.6 mm. At least 8 VI images were quantified from each animal. The median eminence was defined as the base of the mediobasal hypothalamus (far ventral part of the brain), adjacent to the third ventricle from bregma −1.95 to −2.5 mm. Four ME images were quantified from each animal. The area postrema was defined as the region in from bregma −7.2 to −7.75 mm. Four area postrema images were quantified from each animal.

### Microglia morphology analysis

Microglia activation in the hippocampus was quantified using Fiji (ImageJ, NIH). Five images of the dentate gyrus were acquired from each animal. Images were 2048 x 2048 pixels, with a pixel size of 0.315 μm. Images were uploaded to Fiji by a blinded reviewer (KGB) and converted to 8-bit greyscale images. After thresholding, microglia were identified using the “analyze particle” function, which measured mean Iba-1 fluorescent intensity per cell, cell area, and percent area covered by Iba-1 staining.

### *In situ* hybridization

At 10 d.p.i., mice were euthanized with CO_2_ and brains were removed then frozen on dry ice. 20 µm coronal sections were cut on a cryostat and thaw-mounted onto Superfrost Plus slides (VWR Scientific). Sections were collected in a 1:6 series from the diagonal band of Broca (bregma 0.50 mm) caudally through the mammillary bodies (bregma 5.00 mm). 0.15 pmol/ml of an antisense ^33^P-labeled mouse *Ccl2* riboprobe (corresponding to bases 38-447 of mouse *Ccl2*; GenBank accession no. NM_011333.3) was denatured, dissolved in hybridization buffer along with 1.7 mg/ ml tRNA, and applied to slides. Slides were covered with glass coverslips, placed in a humid chamber, and incubated overnight at 55°C. The following day, slides were treated with RNase A and washed under conditions of increasing stringency. Slides were dipped in 100% ethanol, air dried, and then dipped in NTB-2 liquid emulsion (Kodak). Slides were developed 4 d later and cover slipped.

### Quantitative Real-Time PCR

Prior to tissue extraction, mice were euthanized with a lethal dose of a ketamine/xylazine/acetapromide and sacrificed. Hippocampal blocks and gastrocnemii were dissected, snap frozen, and stored in −80 °C until analysis. RNA was extracted using an RNeasy mini kit (Qiagen) according to the manufacturer’s instructions. cDNA was transcribed using TaqMan reverse transcription reagents and random hexamers according to the manufacturer’s instructions. PCR reactions were run on an ABI 7300 (Applied Biosystems), using TaqMan universal PCR master mix with the following TaqMan mouse gene expression assays: *18s* (Mm04277571_s1), *Ccl2* (Mm99999056_m1), *Ccl3* (Mm00441259_g1), *Ccl5* (Mm01302427_m1), *Cxcl1* (Mm04207460_m1), *Cxcl2* (Mm00436450_m1), *Cxcl5* (Mm00436451_g1), *Cxcl9* (Mm00434946_m1), *Cxcl10* (Mm00445235_m1), *Gapdh* (Mm99999915_g1), *Mafbx* (Mm00499518_m1), *Murf1* (Mm01185221_m1), and *Foxo1* (Mm00490672_m1).

Relative expression was calculated using the ΔΔCt method and normalized to WT vehicle treated or sham control. Statistical analysis was performed on the normally distributed ΔCt values.

### Flow cytometry

Mice were anesthetized using a ketamine/xylazine/acetapromide cocktail and perfused with 15 mL ice cold 0.01 M PBS to remove circulating leukocytes. If circulating leukocytes were analyzed, blood was drawn prior to perfusion via cardiac puncture using a 25-gauge needle, then placed in an EDTA coated tube. After perfusion, organs were extracted and immune cells were isolated using the following protocols:

#### Brain

Brains were minced in a digestion solution containing 1 mg/mL type II collagenase (Sigma) and 1% DNAse (Sigma) in RPMI, then placed in a 37°C incubator for 45 min. After digestion, myelin was removed via using 30% percoll in RPMI. Isolated cells were washed with RPMI, incubated in Fc block for 5 min, then incubated in 100 μL of PBS containing antibodies for 30 min at 4°C. Cells were then washed once with RPMI.

#### Liver

Livers were pushed through a 70 μm nylon strainer, then washed once with RPMI. The resulting suspension was resuspended in a 40 mL digestion solution containing 1 mg/mL type II collagenase (Sigma) and 1% DNAse (Sigma) in RPMI, then placed in a 37°C incubator for 1 hr. After digestion, the suspension was placed on ice for 5 min, then the top 35 mL was discarded. The remaining 5 mL was washed in RPMI, resuspended in 10 mL 35% percoll to remove debris, then treated with RBC lysis buffer. The resulting cell suspension was washed with RPMI, then cells were incubated in 100 μL of PBS containing antibodies for 30 min, then washed with RPMI.

#### Tumor

A 0.4-0.5 g piece of pancreatic tumor was removed, minced in a digestion solution containing 1 mg/mL type II collagenase (Sigma) and 1% DNAse (Sigma) in RPMI, then placed in a 37°C incubator for 1 hr. After digestion, cells were washed with RPMI, then incubated in 100 μL of PBS containing antibodies for 30 min at 4°C. Cells were then washed once with RPMI.

#### Blood

200 μL of blood was drawn via cardiac puncture with a 25-gauge needle. Red blood cells were then lysed with 1X RBC lysis buffer. The resulting cell suspension was washed with RPMI, then cells were incubated in 100 μL of PBS containing antibodies for 30 min at 4°C, then washed with RPMI.

#### Gating Strategy

Cells were gated on LD, SSC singlet, and FSC singlet. Immune cells were defined as CD45+ cells. In the brain, microglia were defined as CD45^mid^CD11b+. Leukocytes were identified as either myeloid cells (CD45^high^CD11b+ in the brain, CD45+CD11b+ in all other tissues) or lymphocytes (CD45^high^CD11b-in the brain, CD45+CD11b-in all other tissues). From myeloid cells Ly6C^low^ monocytes (Ly6C^low^Ly6G-), Ly6C^high^ monocytes (Ly6C^high^Ly6G-), and neutrophils (Ly6C^mid^Ly6G+) were identified. From lymphocytes, CD3+ cells were identified as T-cells, and further phenotyped as CD4+ or CD8+ T-cells. CD3-T-cells were divided into NK1.1+ NK cells or CD19+ B-cells. Flow cytometry analysis was performed on a BD Fortessa or LSRII analytic flow cytometer.

#### Antibodies

All antibodies were purchased from BioLegend, except for Live/Dead, which was purchased from Invitrogen (Fixable Aqua, used at 1:200 dilution). The following anti-mouse antibodies were used, with clone, fluorophore, and dilution indicated in parenthesis: CD3 (17A2, PE, 1:100), CD3 (17A2, APC/Cy7, 1:400), CD4 (RM4-5, APC, 1:100), CD8 (53-6.7, APC/Cy7, 1:800), CD11b (M1/70, APC, 1:800), CD11b (M1/70, FITC, 1:200), CD19 (6D5,BV650, 1:33), CD45 (30-F11, PerCP/Cy5.5, 1:400), CD45 (30-F11, APC/Cy7, 1:400), Ly6C (HK1.4, PerCP, 1:100), Ly6G (1A8, PE/Cy7, 1:800), NK1.1 (PK136, BV785, 1:800), CCR2 (SA203G11, AF647, 1:100).

#### FACS sorting for RNAseq

At 10 d.p.i., mice were anesthetized and 200 μL of blood was drawn via cardiac puncture with a 25-gauge needle. Circulating leukocytes were then removed via transcardiac perfusion with PBS and brain, liver, and tumor were removed. Leukocytes were isolated from blood, brain, liver, and tumor as described above. Sorting was performed using an Influx sorter (BD) with a 100 μm nozzle. Neutrophils were defined as CD11b^high^Ly6G^high^ live, singlet cells, and were sorted into lysis buffer (Qiagen), then stored at −80° C.

### RNA Isolation and Sequencing

#### RNA Isolation, sequencing, and library preparation

Total RNA was isolated from FACS-sorted CD11b^high^Ly6G^high^ neutrophils using an RNAeasy Plus Micro kit (Qiagen). RNA integrity was verified by a Bioanalyzer (Agilent). Sample cDNAs were prepared using the SMART-Seq v4 Ultra Low Input kit (Takara) using 250 pg of input total RNA followed by library preparation using a TruSeq DNA Nano kit (Illumina). Libraries were verified by Tapestation (Agilent). Library concentrations were determined by real-time PCR with a StepOnePlus Real Time PCR System (Thermo Fisher) and a Kapa Library Quantification Kit (Kapa Biosystems / Roche). Libraries were sequenced with a 100 cycle single read protocol on a HiSeq 2500 (Illumina) with four libraries per lane. Fastq files were assembled using Bcl2Fastq2 (Illumina).

#### RNA-seq processing and analysis

Quality control checks were done using the FastQC package (https://www.bioinformatics.babraham.ac.uk/projects/fastqc/). Raw reads were normalized and analyzed using the Bioconductor package DESeq2 (Love, Huber, & Anders, 2014), which uses negative binomial generalized linear models. Only those genes that were expressed in at least one sample were included in differential expression analysis. To identify transcripts differentially expressed in brain-infiltrating neutrophils compared to neutrophils infiltrating other organs, gene expression in neutrophils isolated from brain was compared to that in neutrophils isolated from liver, tumor, and blood. In order to control for the effects of tumor on circulating neutrophils, genes that also were differentially expressed in circulating neutrophils from tumor mice compared to circulating neutrophils from sham mice were excluded from analysis. All p-values were adjusted for multiple comparisons using the Benjamani-Hochberg method (Benjamini & Hochberg, 1995). Differential expression was defined based on statistical significance (adjusted p-value < 0.05) and effect size (log_2_ fold change) ≤ or ≥ −2. Heatmaps were created using the pheatmap package from R. Gene Ontology analysis was performed using the Goseq Bioconductor R package(Young, Wakefield, Smyth, & Oshlack, 2010). For pathway enrichment analysis, pathway annotation from the Reactome knowledgebase (Croft et al., 2014; Fabregat et al., 2018) was used.

### Statistical Analysis

Data are expressed as means ± SEM. Statistical analysis was performed with Prism 7.0 software (Graphpad Software Corp). When two groups were compared, data were analyzed with either student’s t-test or Mann-Whitney U test. When more than two groups were compared, data were analyzed with either One-way (when multiple groups were compared to a single sham group) or Two-way (when there were multiple genotypes within tumor and sham groups being compared) ANOVA analysis. For single time point experiments, the two factors in ANOVA analysis were genotype or treatment. In repeated measures experiments, the two factors were group and time. Main effects of genotype, treatment, group, and/or time were first analyzed, and if one effect was significant, Bonferroni *post hoc* analysis was then performed. For all analyses, significance was assigned at the level of *p* < 0.05.

## Acknowledgements

We would like to thank Pamela Canaday, Dorian LaTocha, and Sara Christensen at the OHSU Flow Cytometry Core for their assistance with FACS sorting for RNASeq; Taylor McFarland at the OHSU Gene Profiling Shared Resource for his assistance with RNA isolation; Amy Carlos and Dr. Robert Searles at the OHSU Massively Parallel Sequencing Core for their assistance with RNA sequencing and library preparation; Matthew Michaelis for assistance with the graphical abstract; Dr. Elizabeth Jaffee from Johns Hopkins for providing KPC tumor cells; Drs. Joachim Baumgart and Daniela Reese at Medac GmbH, Germany for providing treosulfan; the Brenden-Colson Center for Pancreatic Care at OHSU for providing funding; and Drs. David Jacoby (OHSU) and Vickie Baracos (University of Alberta) for their help with editing the manuscript. Bioinformatics analytical expertise for this project was supported by partnership between the office of the OHSU Senior Vice President for Research, the University Shared Resources program, the OCTRI Translational Bioinformatics Program (*NIH/NCATS CTSA UL1TR002369)*, and the Integrated Genomics Laboratory’s Massively Parallel Sequencing Shared Resources. This work was supported by National Institutes of Health grants R01CA184324-01 and R01CA217989-01 to D.L. Marks.

## Author Contributions

K.G. Burfeind, D.L. Marks, and K.A. Michaelis designed the research. K.G. Burfeind, M.A. Norgard, X. Zhu, P.R. Levasseur, B. Olson, E.R.S. Torres, and E.M. Patel performed the experiments. K.G. Burfeind, S. Jeng, and S. McWeeney analyzed the data. K.G. Burfeind wrote the manuscript. All authors edited the manuscript. D.L. Marks supervised the study.

## Supplementary Materials

**Figure S1.**
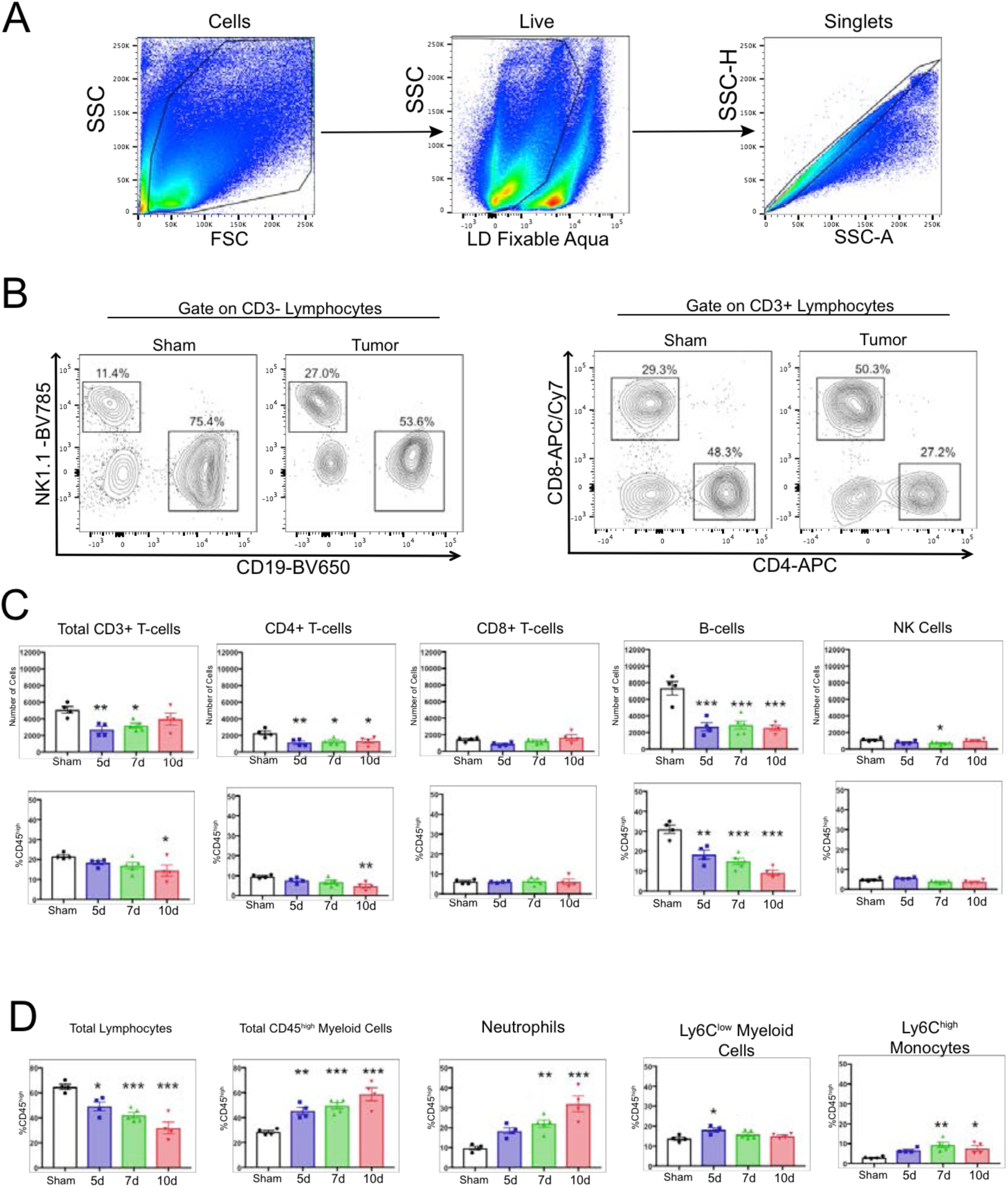
Decreased lymphocytes in the brain during PDAC. A) Gating strategy to identify live single cells from whole brain homogenate. B) Representative plots of different lymphocyte populations from brain homogenate from sham and tumor (10 d.p.i.) animals. For CD3- cells, NK cells = NK1.1+CD19-, B-cells = CD19+NK1.1-. For CD3+ cells, CD4+ and CD8+ T-cells were identified. C) Quantification of different lymphocyte populations throughout PDAC. \**P* < 0.05, \*\**P* < 0.01, \*\*\**P* < 0.001 compared to sham one-way ANOVA Bonferroni *post hoc* analysis. D) Quantification of different immune cell populations in the brain, as a percentage of CD45^high^ cells. \**P* < 0.05, \*\**P* < 0.01, \*\*\**P* < 0.001 compared to sham. *n* = 4-5/group. Results are representative of three independent experiments.

**Figure S2.**
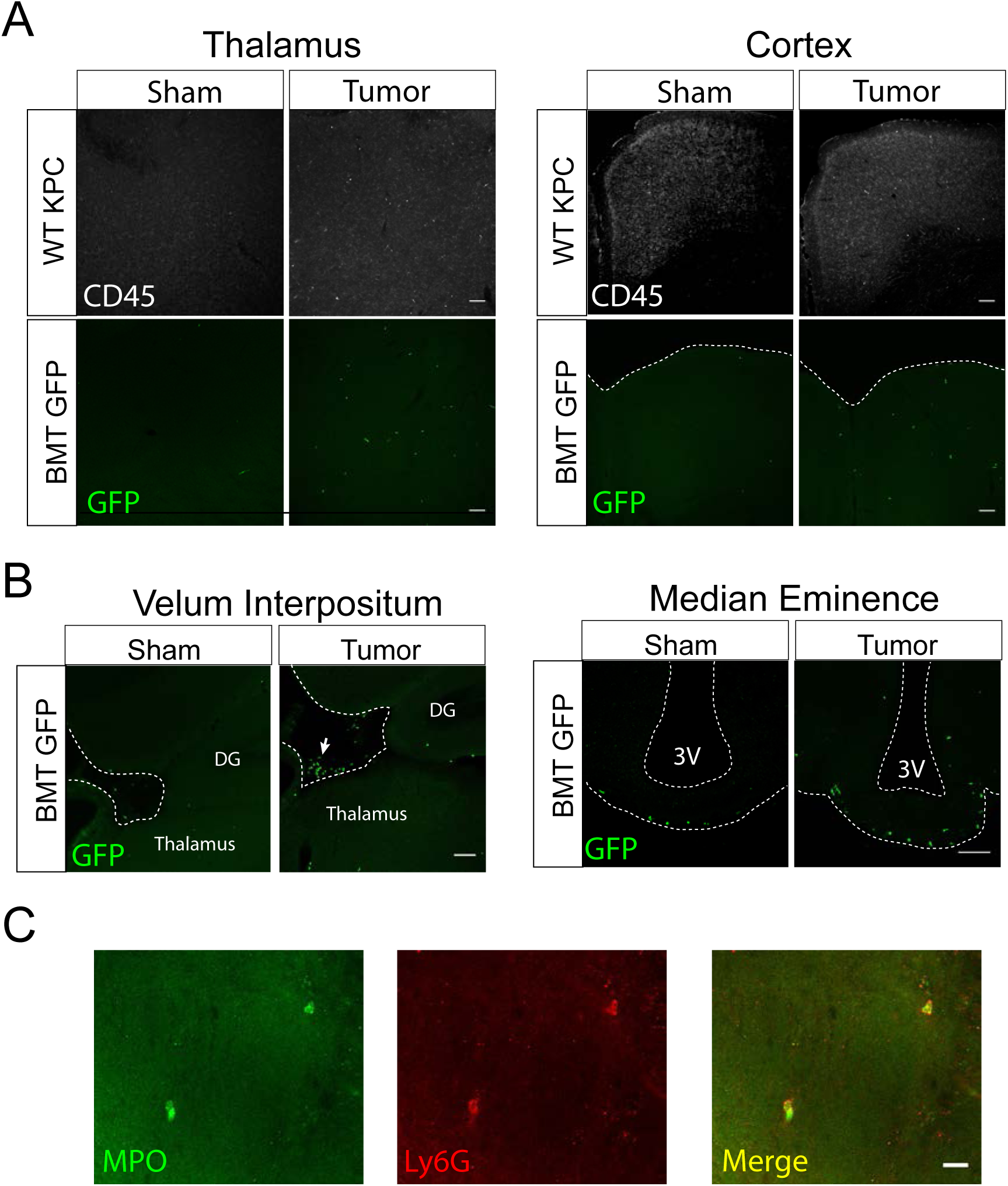
Immunofluorescence analysis of infiltrating immune cells during PDAC. A) 10X confocal images of thalamus and cortex from sham and tumor mouse brains, 10 d.p.i. WT KPC = WT animals, BMT GFP = Ly5.1 eGFP marrow transplanted into WT recipient after treosulfan conditioning to ablate marrow (see Methods). Scale bar = 100 μm. B) 10X (VI) and 20X (ME) confocal images of VI and ME from sham and tumor (10 d.p.i.) mice. In images of the VI, dashed line denotes edge of parenchyma and beginning of meninges. Arrow = cluster of infiltrating GFP+ immune cells in the VI meninges. DG = dentate gyrus. 3V = third ventricle. Scale bars = 100 μm. C) 40X confocal image of thalamus from tumor mouse, 12 d.p.i. Scale bar = 20 μm.

**Figure S3:**
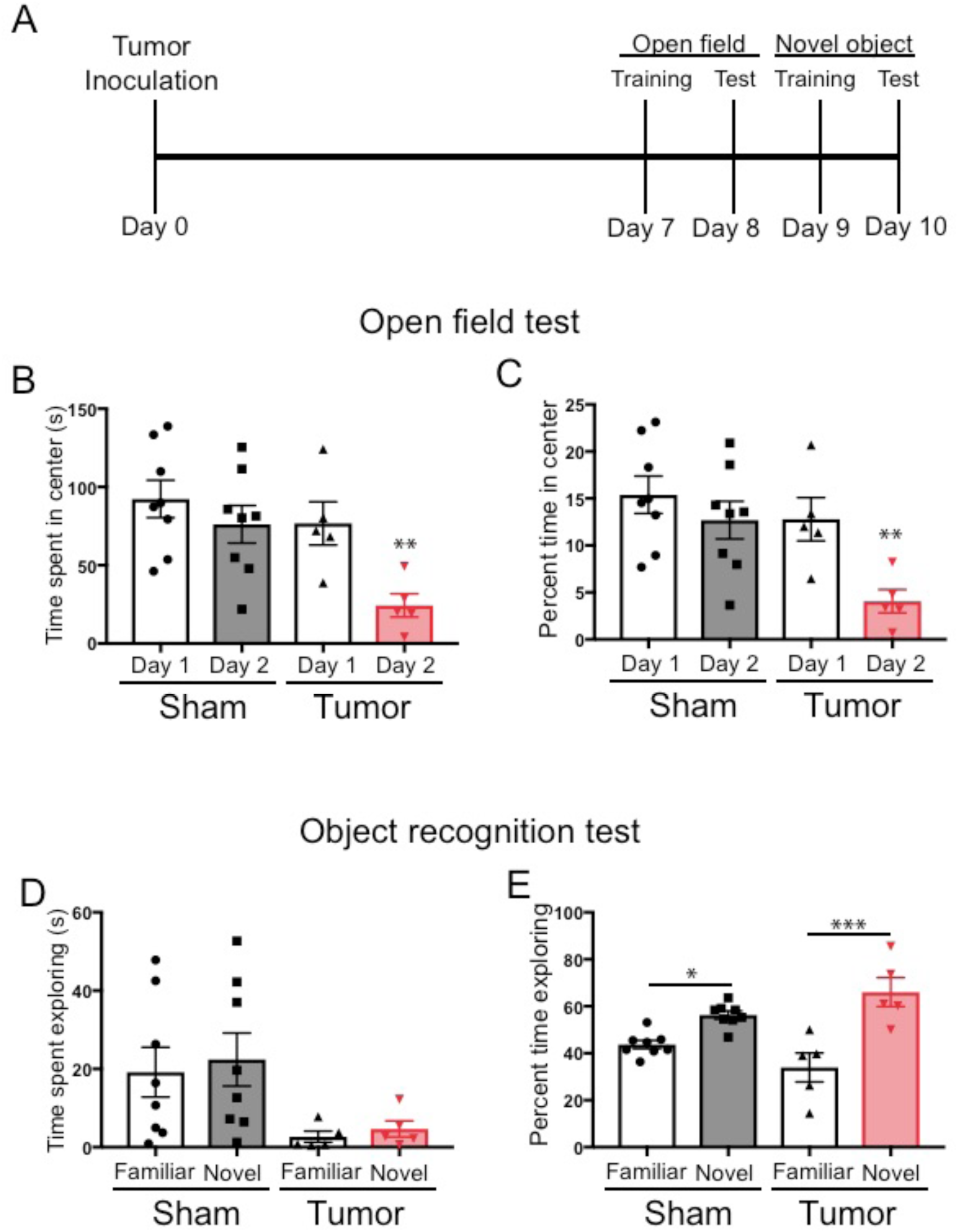
Cognitive performance of KPC tumor mice. A) Illustration depicting timeline of behavioral tests. B) Total time spent in center of arena during open field test on the first and second day of testing. C) Percent of total time that was spent in the center of the arena during the first and second day of the open field test. For B and D, \*\**P* < 0.01, student’s t-test comparing sham day 2 to tumor day 2. D) Total time spent investigating the familiar or the novel object. E) Percent of total time spent exploring the familiar object and the novel object. *\*P* < 0.05, \*\*\**P* < 0.001, Bonferroni post-hoc analysis in two-way ANOVA. For all panels, *n* = 8 sham mice and *n* = 5 tumor mice. Three tumor animals were excluded from all analyses due to complete lack of movement. Data are presented as mean ± s.e.m.

**Figure S4.**
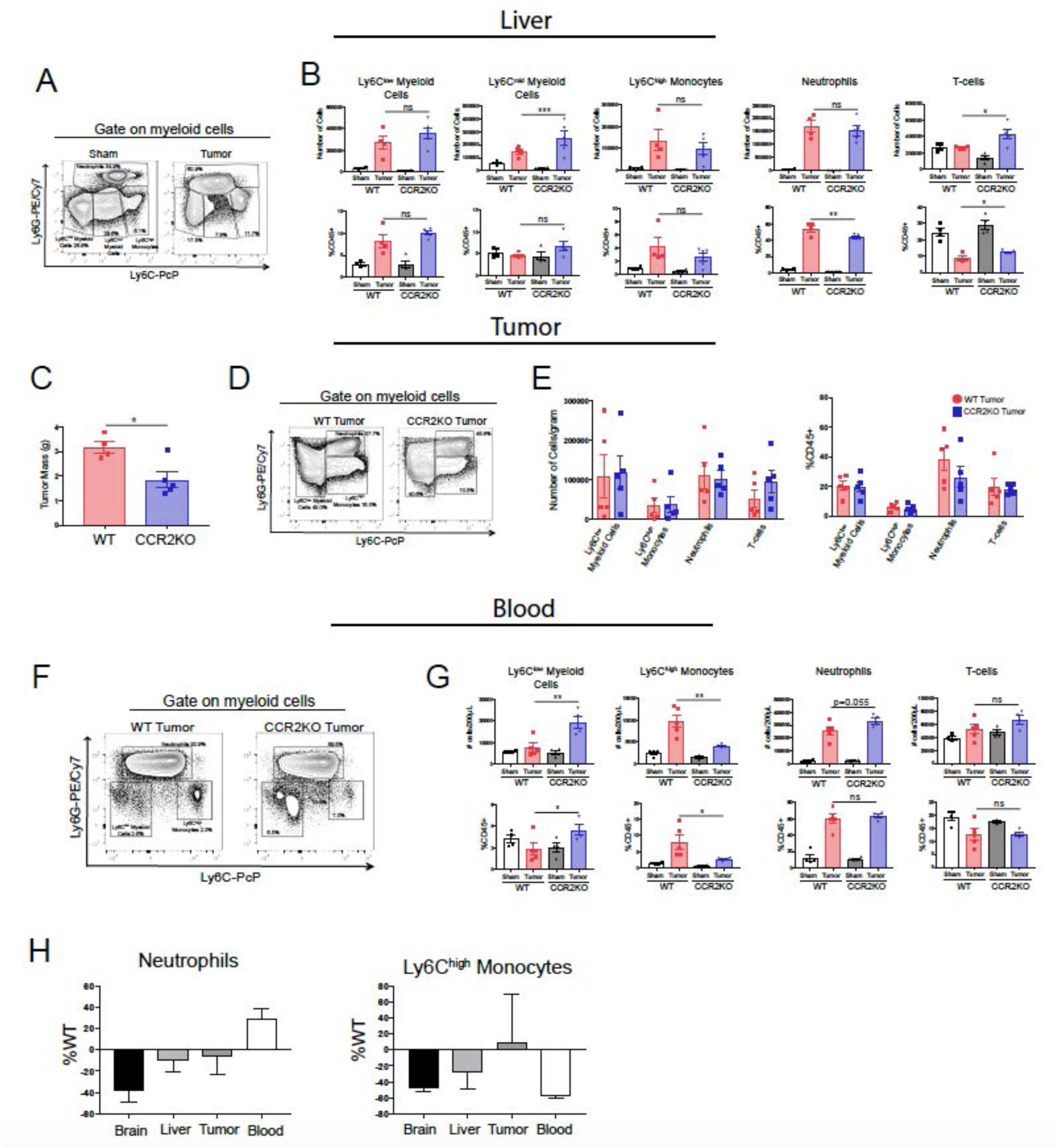
The CCR2-CCL2 axis is of selective importance for the brain during PDAC. A) Representative flow cytometry plot of different myeloid cell populations from WT sham and tumor livers, 11 d.p.i., in order to illustrate different myeloid cell populations identified based on Ly6C and Ly6G expression. Cells are gated on live, singlet CD45+CD11b+ cells. B) Quantification of flow cytometry analysis of different immune cell populations in the liver from WT and CCR2KO sham and tumor animals, 11 d.p.i. \**P* < 0.05, \*\**P* < 0.01, WT tumor vs. CCR2KO tumor, or tumor vs. sham in the same genotype in Bonferroni *post hoc* analysis in two-way ANOVA. ns = not significant. *n* = 4-9/group. C) Tumor mass from WT and CCR2KO animals, 11 d.p.i. Data are representative of three independent experiments. Data are presented as mean ± s.e.m. D) Representative flow cytometry plot of different myeloid cell populations from WT and CCR2KO tumors, 10 d.p.i. Cells are gated on live, singlet CD45+CD11b+ cells. E) Quantification of flow cytometry analysis of different immune cell populations isolated from tumor from WT and CCR2KO tumor animals, 10 d.p.i. Data consist of two independent experiments pooled (*n* = at least 2 per group per experiment). Data are presented as mean ± s.e.m. F) Representative plot of different myeloid cell populations from WT and CCR2KO tumor animal blood, 10 d.p.i. Cells are gated on live, singlet CD45+CD11b+ cells. G) Quantification of flow cytometry analysis of different immune cell populations in the blood from WT and CCR2KO sham and tumor animals, 10 d.p.i. \**P* < 0.05, \*\**P* < 0.01, WT tumor vs. CCR2KO tumor, or tumor vs. sham in the same genotype in Bonferroni *post hoc* analysis in two-way ANOVA. ns = not significant. *n* =4-5/group. Data are representative of two independent experiments. H) Analysis of neutrophils and Ly6C^high^ monocytes in brain, liver, tumor, and blood in CCR2KO tumor mice, normalized to number in WT tumor mice. *n* = 5-9/group.

**Figure S5.**
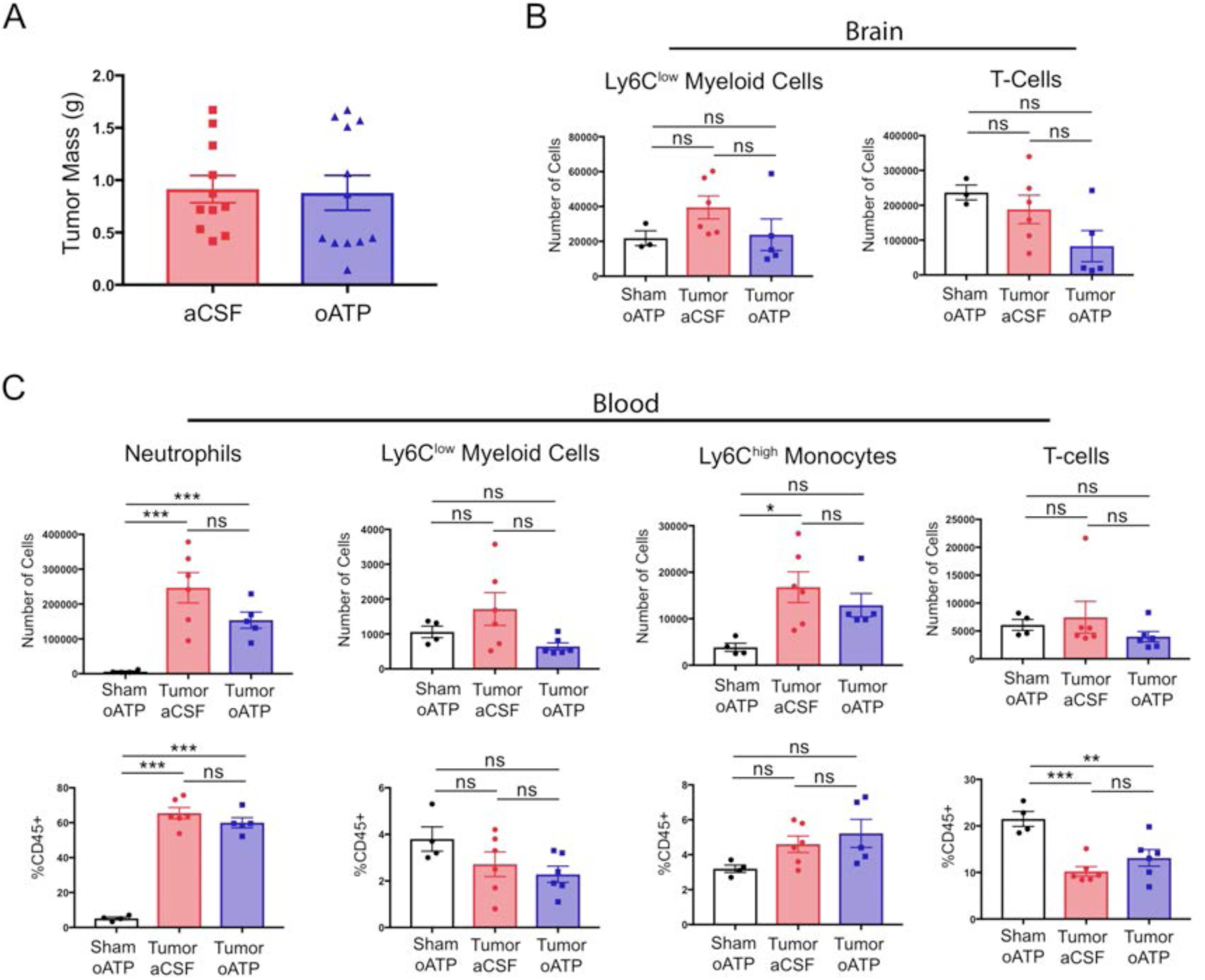
Intracerebroventricular antagonism of P2RX7 does not affect systemic inflammation or tumor size during PDAC. A) Tumor mass from aCSF- and oATP-treated tumor-bearing mice, 8-10 d.p.i. *n* = 11-12/group. Results consist of two independent experiments pooled (*n* = 5-7/group in each experiment). B) Quantification of immune cells isolated from whole brain homogenate. ns = not significant in Bonferroni *post hoc* analysis in two-way ANOVA. *n* = 4-7/group. C) Quantification of immune cells isolated from blood, per 200 µL of blood. \**P* < 0.05, \*\**P* < 0.01, \*\*\**P* < 0.001 in Bonferroni *post hoc* analysis in two-way ANOVA. ns = not significant. *n* = 4-6/group.

**Figure 1 – Figure Supplement 1.**
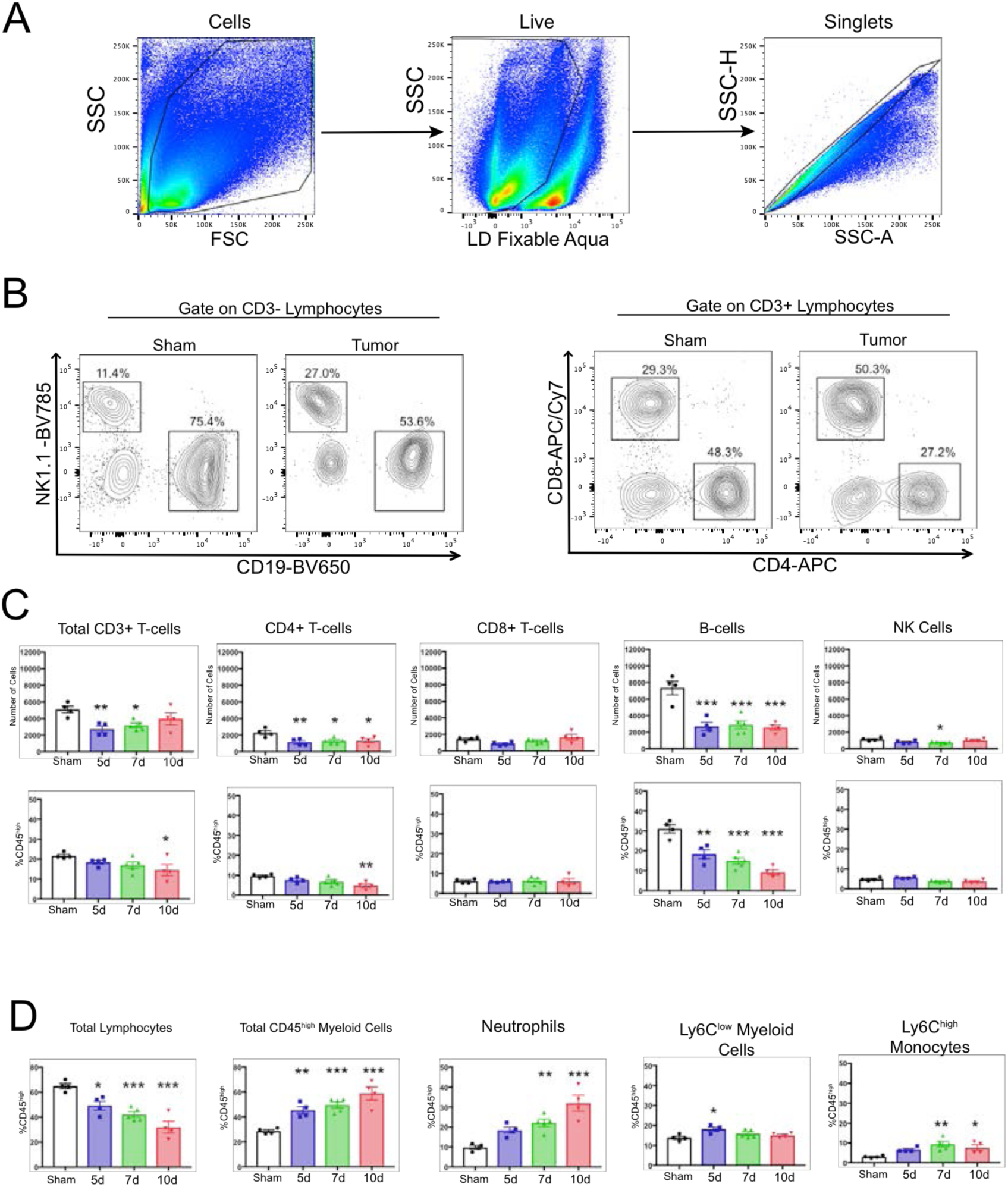
Decreased lymphocytes in the brain during PDAC cachexia. A) Gating strategy to identify live single cells from whole brain homogenate. B) Representative plots of different lymphocyte populations from brain homogenate from sham and tumor (10 d.p.i.) animals. For CD3- cells, NK cells = NK1.1+CD19-, B-cells = CD19+NK1.1-. For CD3+ cells, CD4+ and CD8+ T-cells were identified. C) Quantification of different lymphocyte populations throughout the course of cachexia. \**P* < 0.05, \*\**P* < 0.01, \*\*\**P* < 0.001 compared to sham one-way ANOVA Bonferroni *post hoc* analysis. D) Quantification of different immune cell populations in the brain throughout the course of cachexia, as a percentage of CD45^high^ cells. \**P* < 0.05, \*\**P* < 0.01, \*\*\**P* < 0.001 compared to sham. *n* = 4-5/group. Results are representative of three independent experiments.

**Figure 1 – Figure Supplement 2.**
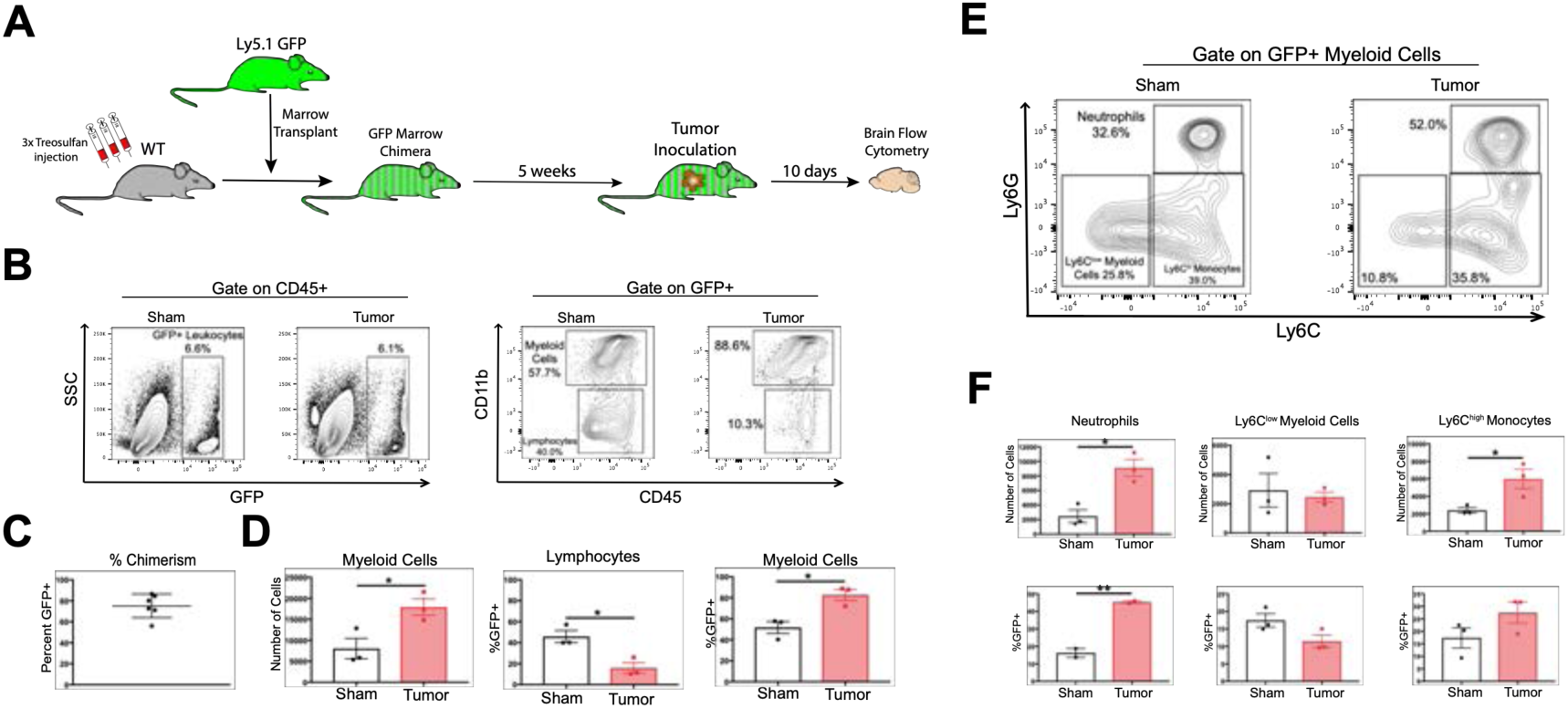
GFP BMT confirms peripheral origin of infiltrating myeloid cells in the CNS during PDAC. A) Diagram of bone marrow transplant protocol to generate GFP+ bone marrow chimeras. B) Gating strategy for CD45+GFP+ cells isolated from brains of tumor and sham GFP chimera animals. C) Percent chimerism, identified as percentage of CD45+ cells in the blood that were GFP+. D) Quantification of GFP+ myeloid cells and lymphocytes in the brains of tumor and sham mice, 10 d.p.i. E) Representative flow cytometry plot of different GFP+ myeloid cell populations in the brains of tumor and sham GFP bone marrow chimera animals, 10 d.p.i. F) Quantification of different GFP+ myeloid cell populations in the brains of tumor and sham GFP bone marrow chimera animals, 10 d.p.i., as identified in Fig. 1k. *n* = 3/group, \**P* < 0.05, \*\**P* < 0.01 in student’s t-test.

**Figure 2 – Figure Supplement 1.**
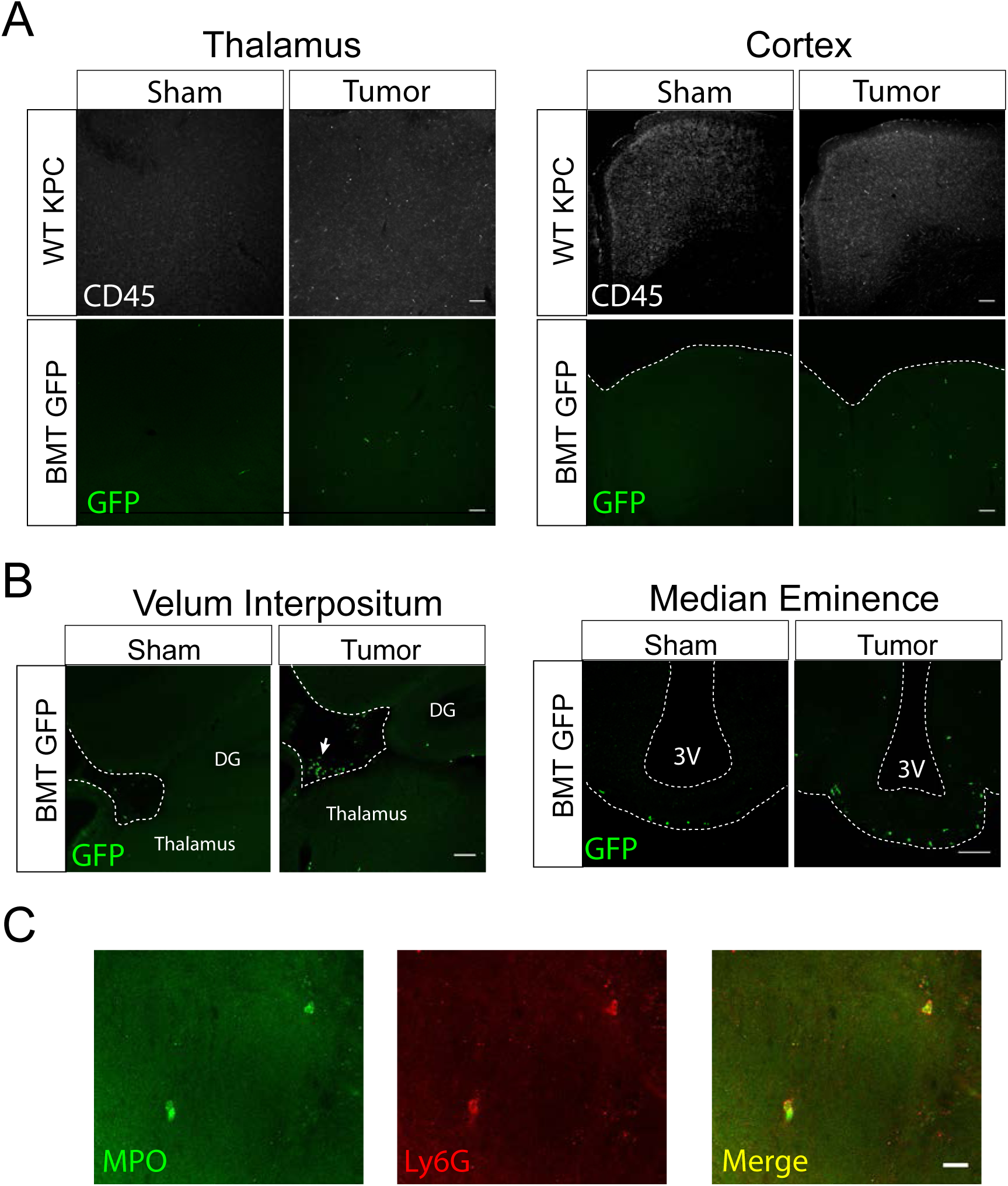
Immunofluorescence analysis of infiltrating immune cells during PDAC. A) 10X confocal images of thalamus and cortex from sham and tumor mouse brains, 10 d.p.i. WT KPC = WT animals, BMT GFP = Ly5.1 eGFP marrow transplanted into WT recipient after treosulfan conditioning to ablate marrow (see Methods). Scale bar = 100 μm. B) 10X (VI) and 20X (ME) confocal images of VI and ME from sham and tumor (10 d.p.i.) mice. In images of the VI, dashed line denotes edge of parenchyma and beginning of meninges. Arrow = cluster of infiltrating GFP+ immune cells in the VI meninges. DG = dentate gyrus. 3V = third ventricle. Scale bars = 100 μm. C) 40X confocal image of thalamus from tumor mouse, 12 d.p.i. Scale bar = 20 μm.

**Figure 2 – Figure Supplement 2.**
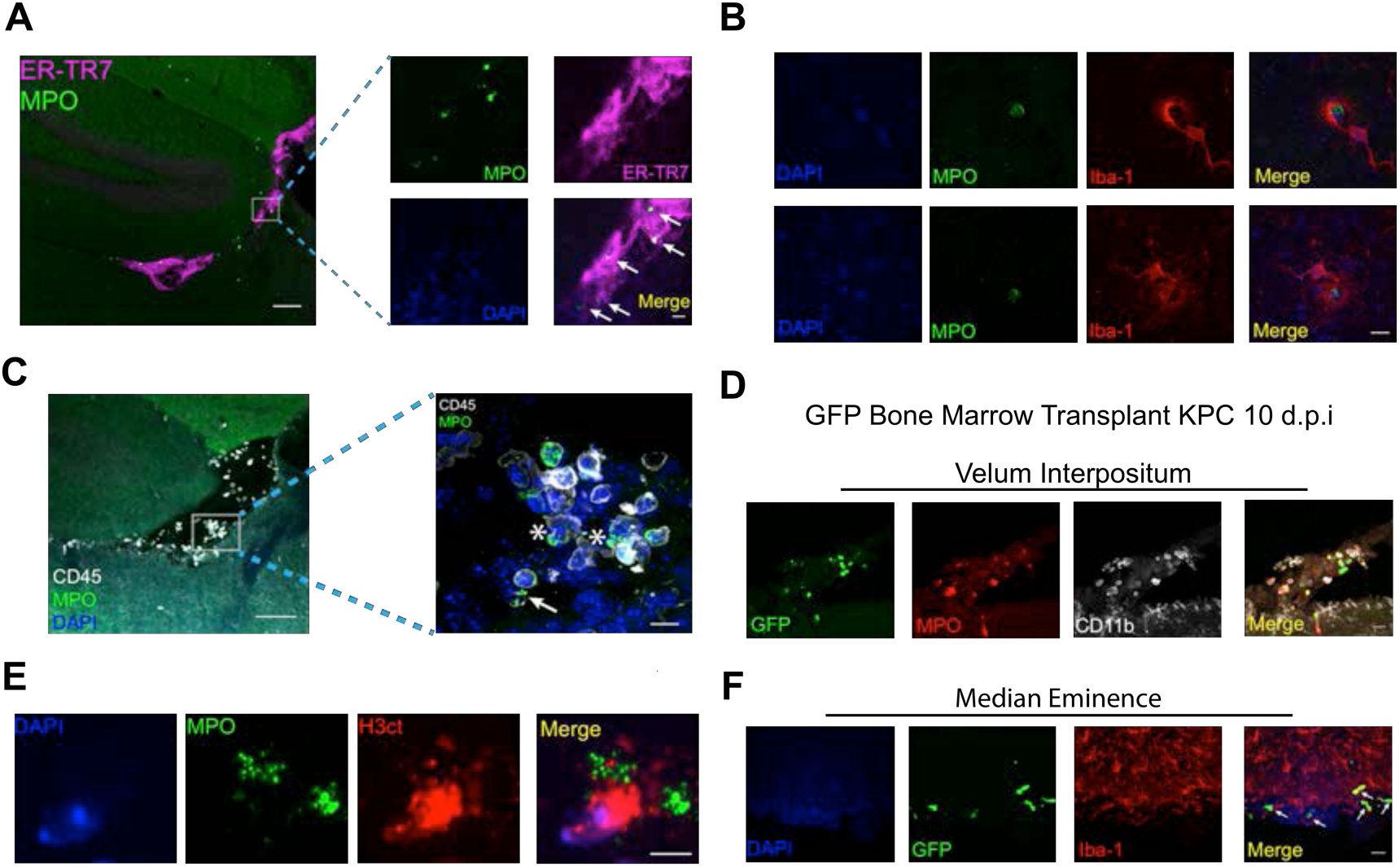
Characteristics of brain-infiltrating immune cells during PDAC. A) Representative 10X image of VI from the brain of a tumor animal 10 d.p.i., showing ER-TR7 staining to label meninges and MPO staining to label neutrophils. Scale bar = 100 μm. Inset = 60X showing neutrophils (indicated by arrows) within the meninges of the VI. Scale bar = 10 μm. B) 20X image a VI with 60X inset showing neutrophils degranulating. Asterisk = myeloperoxidase “blebs” coming off neutrophil. Arrow = extracellular myeloperoxidase. C) 60X image of neutrophil extracellular trap in the VI of a tumor animal, 10 d.p.i. Scale bar = 5 μm. D) Representative 60X images of microglia phagocytosing neutrophils in the thalamus of animals with KPC tumor, 10 d.p.i. Scale bar = 10 μm. E & F) Representative 60X images of VI and ME, respectively, from BMT GFP tumor mice, at 10 d.p.i. Scale bars = 20 μm. Arrows = GFP+Iba-1+ infiltrating macrophages.

**Figure 2 – Figure Supplement 3.**
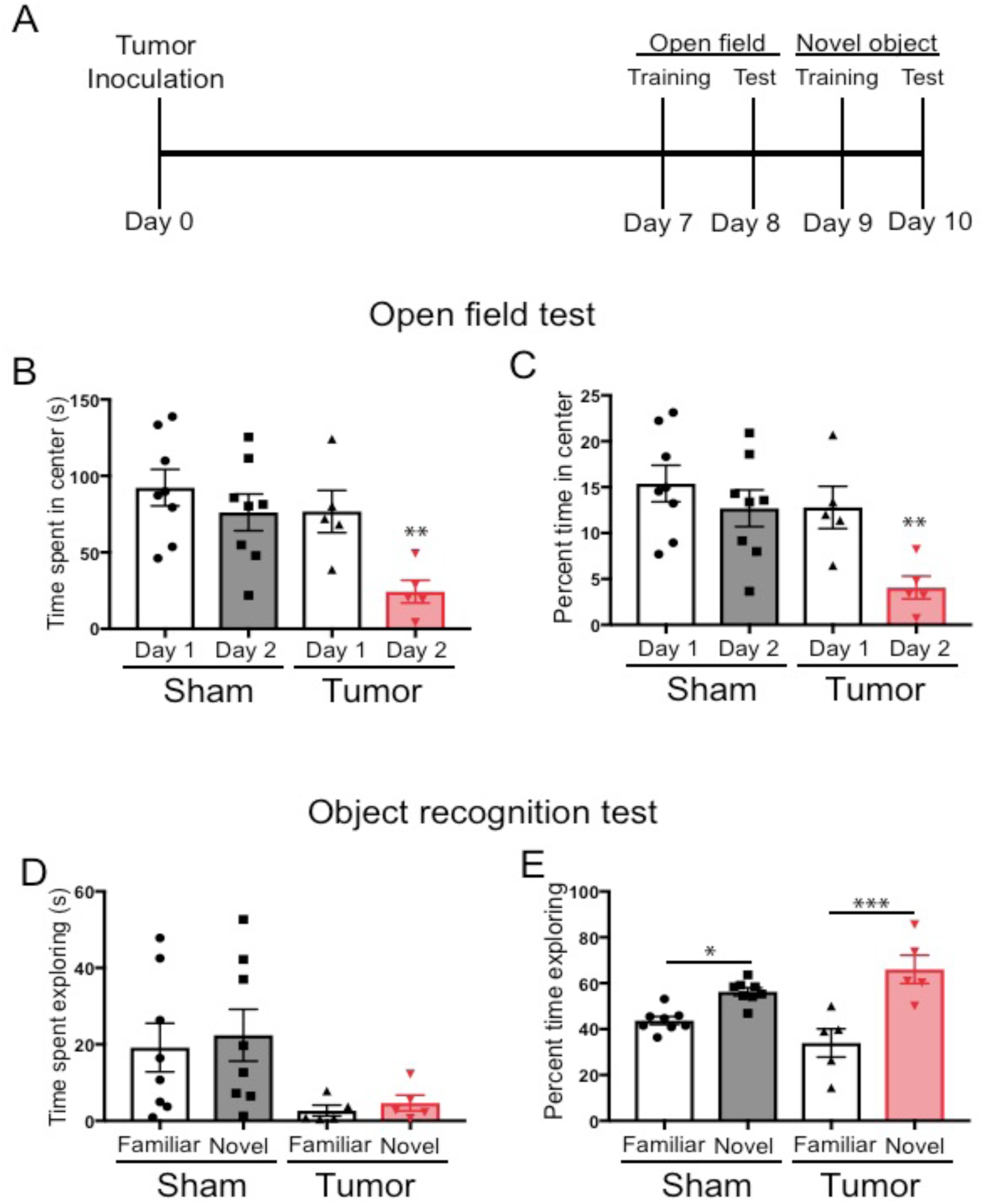
Cognitive performance of KPC tumor mice. A) Illustration depicting timeline of behavioral tests. B) Total time spent in center of arena during open field test on the first and second day of testing. Overall, we observed that tumor mice were significantly less mobile than sham mice in the open field (Fig. S3B and D), which is consistent with our previous home cage locomotor activity analysis showing that KPC tumor mice exhibit decreased activity throughout the course of disease (Michaelis et al., 2017).C) Percent of total time that was spent in the center of the arena during the first and second day of the open field test. For B and D, \*\**P* < 0.01, student’s t-test comparing sham day 2 to tumor day 2. D) Total time spent investigating the familiar or the novel object. E) Percent of total time spent exploring the familiar object and the novel object. *\*P* < 0.05, \*\*\**P* < 0.001, Bonferroni post-hoc analysis in two-way ANOVA. For all panels, *n* = 8 sham mice and *n* = 5 tumor mice. Three tumor animals were excluded from all analyses due to complete lack of movement. Data are presented as mean ± s.e.m.

**Figure 3 – Figure Supplement 1.**
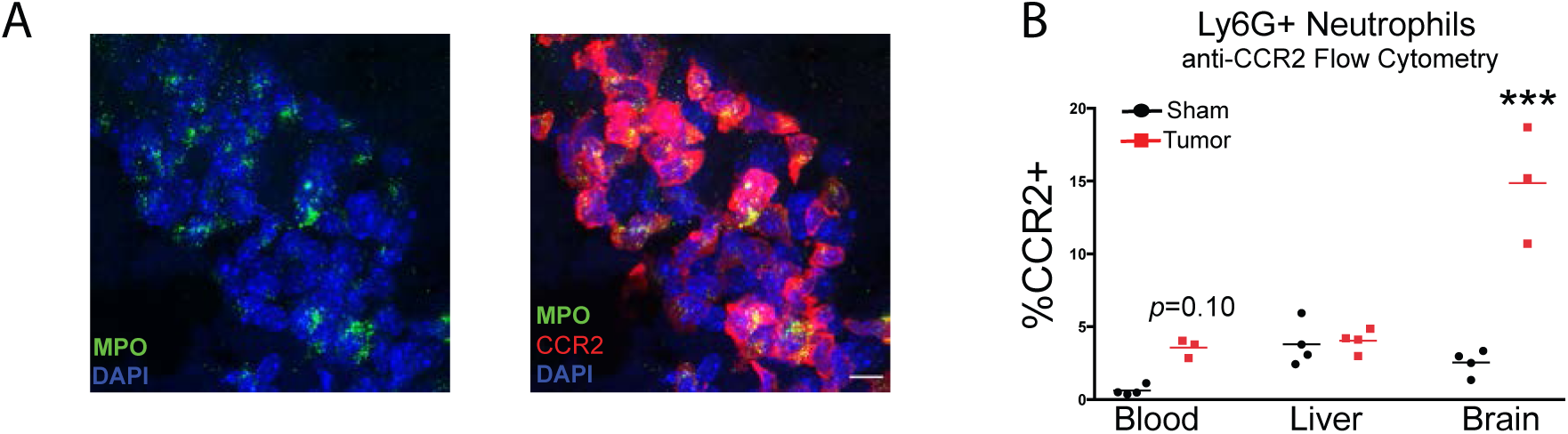
Neutrophils in the Velum Interpositum Express CCR2 during PDAC. A) 60X image identifying cluster of CCR2+ neutrophils in the VI of a tumor animal. Scale bar = 10 μm. B) Flow cytometry analysis of CCR2+ neutrophils in the blood, brain, and liver of sham and tumor animals, 10 d.p.i. Neutrophils defined as live, CD45^high^CD11b+Ly6G+ cells. CCR2+ neutrophils identified by AF647 anti-CCR2 labeling compared to AF647 isotype control. *n* = 3-4/group. *\*\*\*P* < 0.001 in repeated measures one-way ANOVA compared to sham. Bars denote mean. Results are representative of two independent experiments.

**Figure 3 – Figure Supplement 2.**
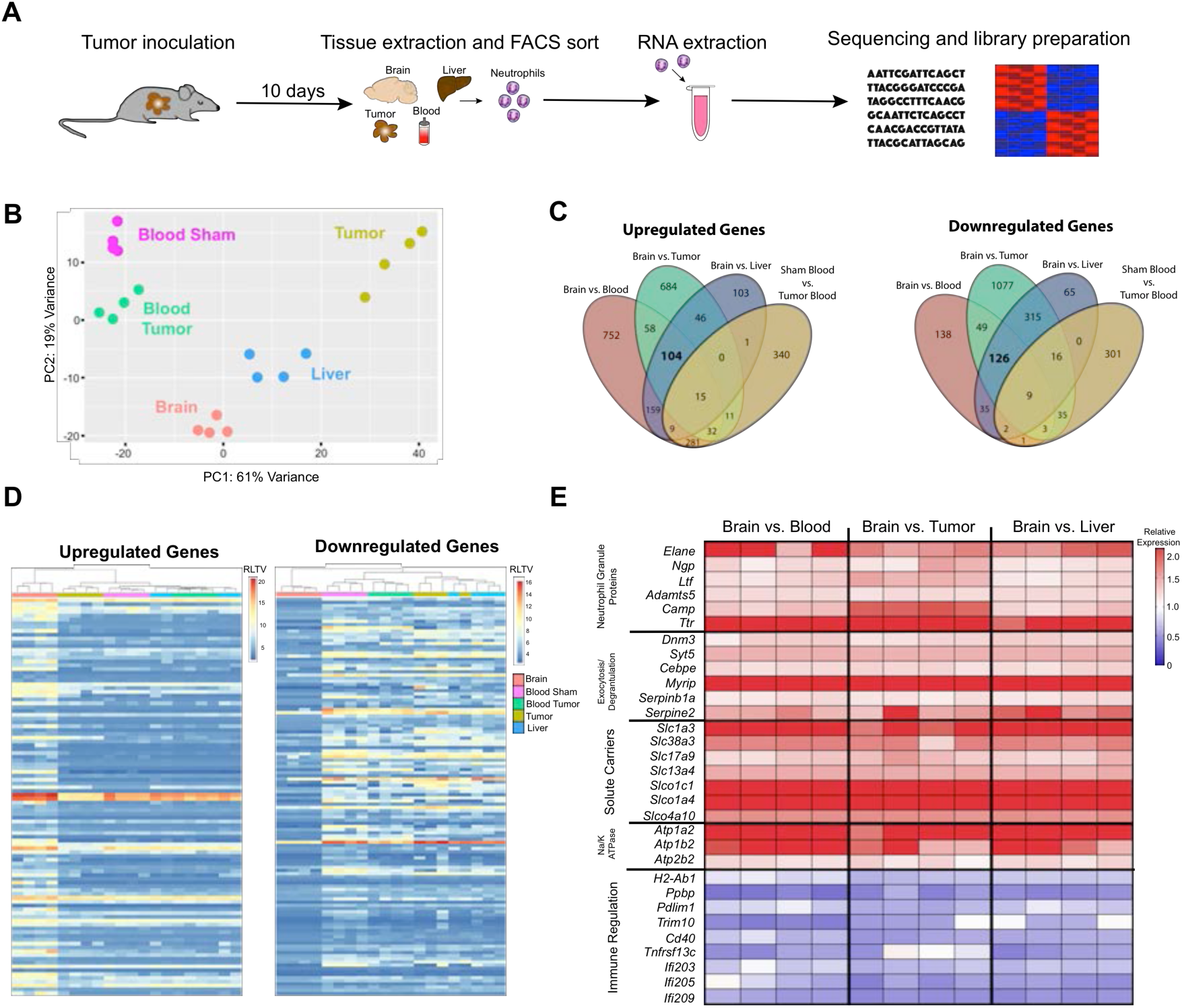
Brain-infiltrating neutrophils express a unique transcriptome during PDAC. A) Workflow for neutrophil isolation, RNA extraction, and RNAseq analysis. B) Principle component analysis of 500 most varying genes in neutrophils isolated from blood, tumor, liver, and brain from mice with PDAC at 10 d.p.i., as well as blood from sham mice. C) Venn diagram of different comparisons of transcripts expressed in neutrophils from different organs. D) We identified putative “brain-specific” transcripts by comparing the transcriptome of brain-infiltrating neutrophils to that of liver- and tumor-infiltrating neutrophils, as well as circulating neutrophils (all from tumor animals). In order to control for the nonspecific effects of malignancy on circulating neutrophils, we any excluded transcripts that were upregulated in circulating neutrophils from tumor animals compared to circulating neutrophils from sham animals. Using this approach, we identified 104 upregulated and 126 downregulated “brain-specific” transcripts RLTV = regularized logarithm transformed value. E) Heatmap of select brain-specific transcripts showing relative expression, comparing average of brain neutrophils to neutrophils in different organs. Functional enrichment analysis (based on Gene Ontology curation) of brain-specific transcripts identified enrichment for the term “extracellular space” (GO:0005615) in upregulated genes and enrichment for the terms “external side of plasma membrane” (GO:0009897), “immune response” (GO:0006955), and “response to interferon-gamma” (GO: 00034341) in downregulated genes. Several brain-specific upregulated transcripts encoded neutrophil granule components and enzymes, such as neutrophil granule protein (*Ngp*), the metalloproteinase ADAMTS5 (*Adamts5*) neutrophil elastase (*Elane*), lactoferrin (*Ltf*), cathelicidin antimicrobial peptide (*Camp*), and transthyretin (*Ttr*), as well as proteins important for granule secretion and NET formation such as dynamin 3 (*Dnm3*), synaptotagmin 15 (*Syt15*), Serpinb1a (*Serpinb1a*), Serpin Family E Member 2 (*Serpine2*), C/EBPε (*Cebpe*) (Gombart et al., 2003), and Myosin VIIA And Rab Interacting Protein (*Myrip*) (Desnos et al., 2003). We also observed an increase in genes for solute carriers (*Slc* gene family) and components of the Na/K ATPase, suggesting brain-infiltrating neutrophils are highly metabolically active. Many brain-specific downregulated transcripts encoded proteins important for immune function and responsiveness to T-cell-derived cytokines, such as MHC II (*H2-Ab1*), CXCL7 (*Ppbp*), PDLIM1 (*Pdlim1*, a negative regulator of NFκβ signaling), and the interferon-inducible genes *Ifi203*, *Ifi205*, and *Ifi209*.

**Figure 3 – Figure Supplement 3.**
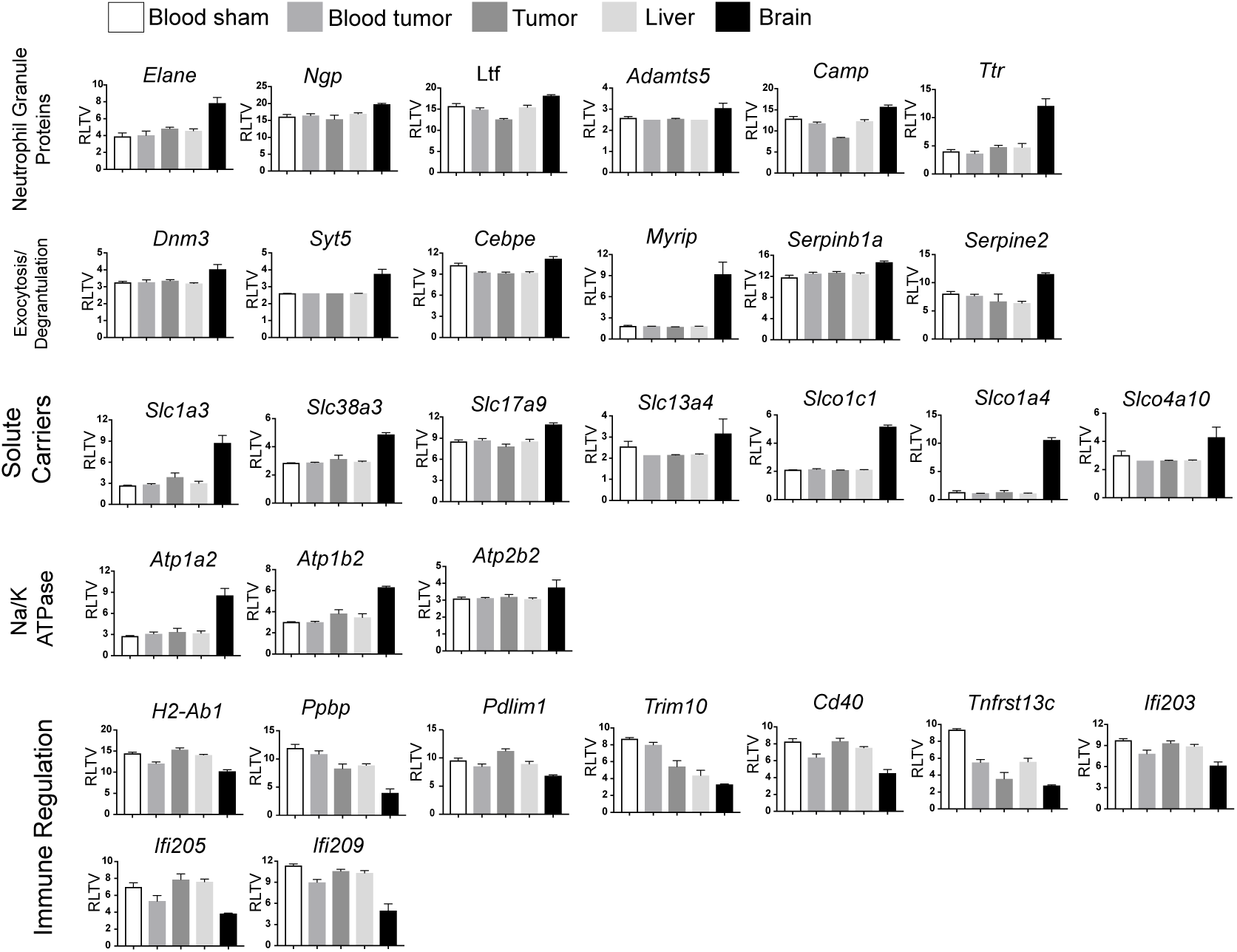
Expression of neutrophil “brain-specific” transcripts. Normalized expression values of transcripts depicted in heatmap in Figure 7e. RLTV = regularized logarithm transformed value.

**Figure 4 – Figure Supplement 1.**
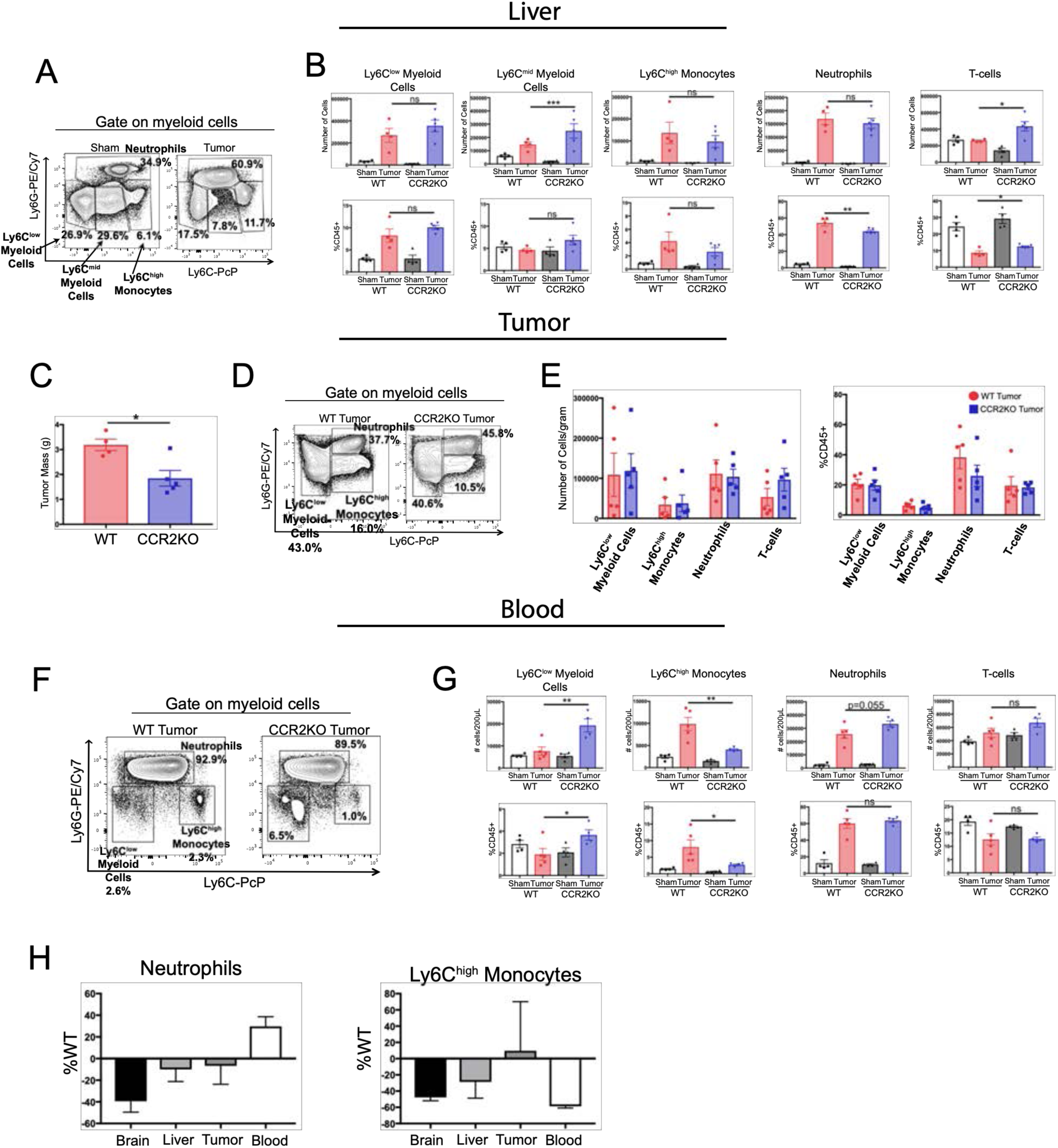
The CCR2-CCL2 axis is of selective importance for the brain in PDAC cachexia. A) Representative flow cytometry plot of different myeloid cell populations from WT sham and tumor livers, 11 d.p.i., in order to illustrate different myeloid cell populations identified based on Ly6C and Ly6G expression. Cells are gated on live, singlet CD45+CD11b+ cells. Quantification of flow cytometry analysis of different immune cell populations in the liver from WT and CCR2KO sham and tumor animals, 11 d.p.i. \**P* < 0.05, \*\**P* < 0.01, WT tumor vs. CCR2KO tumor, or tumor vs. sham in the same genotype in Bonferroni *post hoc* analysis in two-way ANOVA. ns = not significant. *n* = 4-9/group. C) Tumor mass from WT and CCR2KO animals, 11 d.p.i. Data are representative of three independent experiments. Data are presented as mean ± s.e.m. D) Representative flow cytometry plot of different myeloid cell populations from WT and CCR2KO tumors, 10 d.p.i. Cells are gated on live, singlet CD45+CD11b+ cells. E) Quantification of flow cytometry analysis of different immune cell populations isolated from tumor from WT and CCR2KO tumor animals, 10 d.p.i. Data consist of two independent experiments pooled (*n* = at least 2 per group per experiment). Data are presented as mean ± s.e.m. F) Representative plot of different myeloid cell populations from WT and CCR2KO tumor animal blood, 10 d.p.i. Cells are gated on live, singlet CD45+CD11b+ cells. G) Quantification of flow cytometry analysis of different immune cell populations in the blood from WT and CCR2KO sham and tumor animals, 10 d.p.i. \**P* < 0.05, \*\**P* < 0.01, WT tumor vs. CCR2KO tumor, or tumor vs. sham in the same genotype in Bonferroni *post hoc* analysis in two-way ANOVA. ns = not significant. *n* =4-5/group. Data are representative of two independent experiments. H) Analysis of neutrophils and Ly6C^high^ monocytes in brain, liver, tumor, and blood in CCR2KO tumor mice, normalized to number in WT tumor mice. *n* = 5-9/group.

**Figure 5 – Figure Supplement 1.**
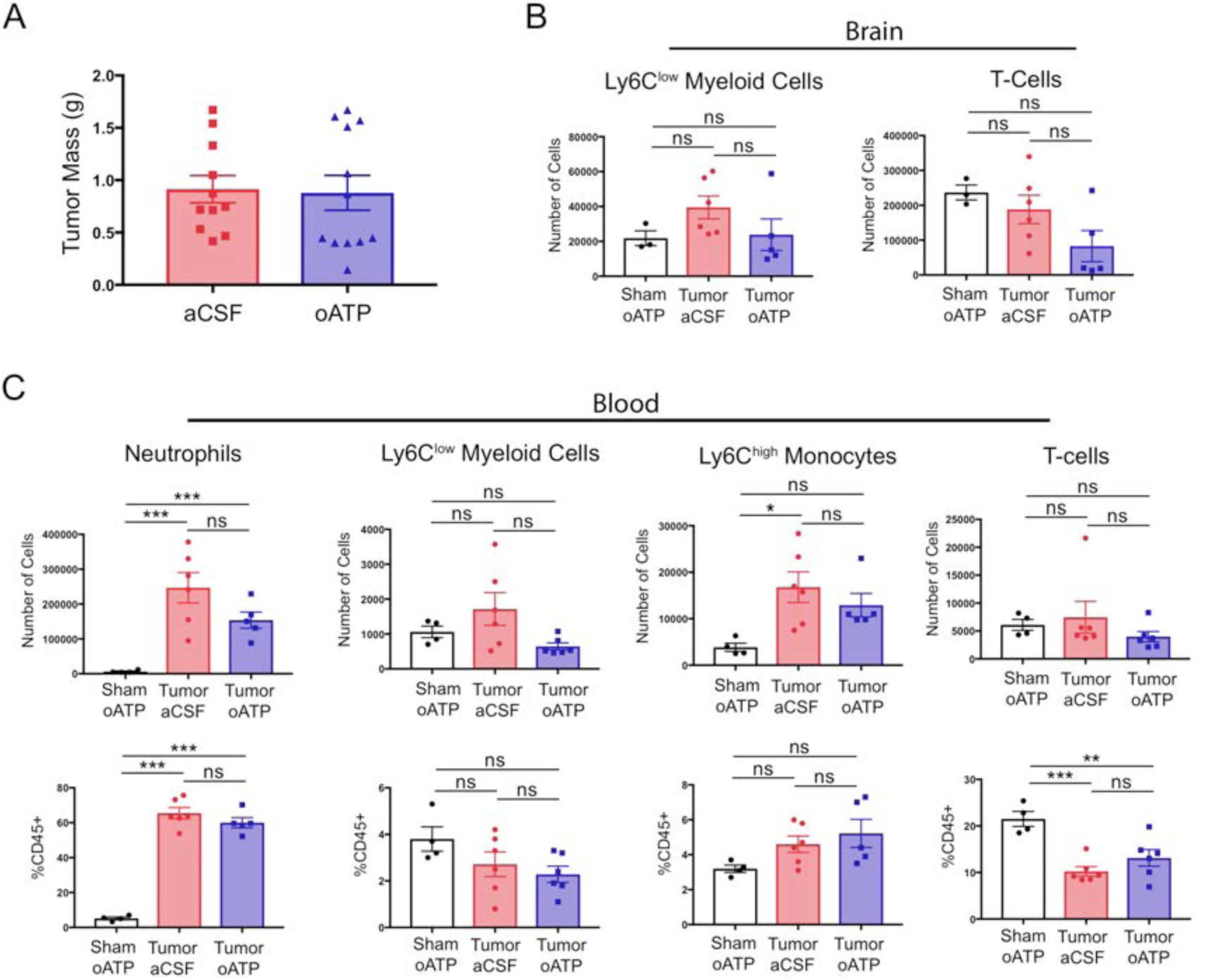
Intracerebroventricular antagonism of P2RX7 does not affect systemic inflammation or tumor size during PDAC. A) Tumor mass from aCSF- and oATP-treated tumor-bearing mice, 8-10 d.p.i. *n* = 11-12/group. Results consist of two independent experiments pooled (*n* = 5-7/group in each experiment). B) Quantification of immune cells isolated from whole brain homogenate. ns = not significant in Bonferroni *post hoc* analysis in two-way ANOVA. *n* = 4-7/group. C) Quantification of immune cells isolated from blood, per 200 µL of blood. \**P* < 0.05, \*\**P* < 0.01, \*\*\**P* < 0.001 in Bonferroni *post hoc* analysis in two-way ANOVA. ns = not significant. *n* = 4-6/group.

**Figure 5 – Figure Supplement 2.**
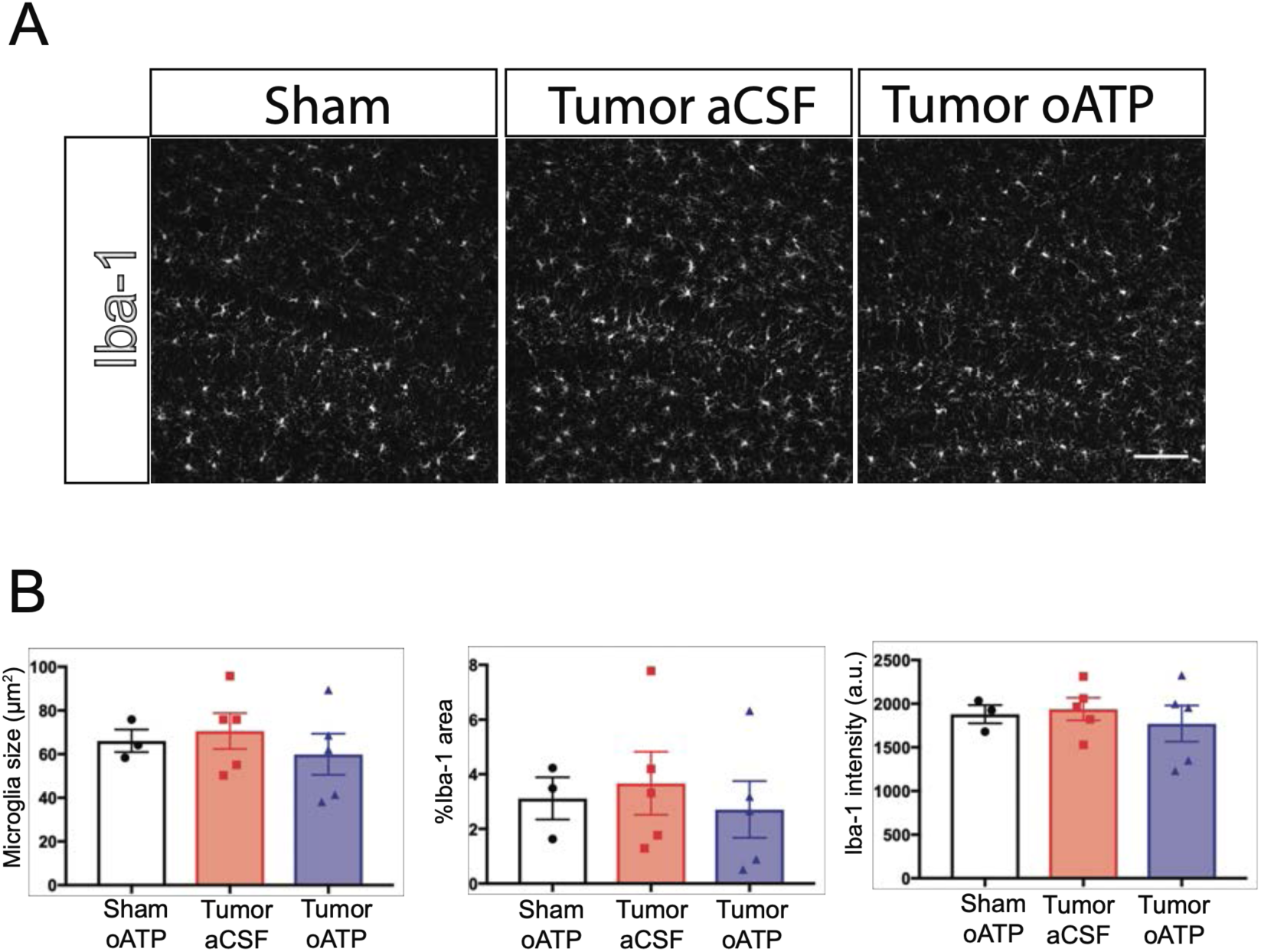
Intracerebroventricular administration of oxidized ATP does not affect microglia activation during PDAC. A) Representative 20X images of Iba-1 immunofluorescence in the dentate gyrus, 10 d.p.i. Scale bar = 100 µm. B) Quantification of microglia morphology in the dentate gyrus 10 d.p.i., showing mean microglia size (left), percent area covered by Iba-1 immunofluorescence (middle), and mean Iba-1 fluorescent intensity per microglia (right). a.u. = arbitrary units. *n* = 3-5/group.

